# Tsr4 is a cytoplasmic chaperone for the ribosomal protein Rps2 in *Saccharomyces cerevisiae*

**DOI:** 10.1101/559146

**Authors:** Joshua J. Black, Sharmishtha Musalgaonkar, Arlen W. Johnson

## Abstract

Eukaryotic ribosome biogenesis requires the action of approximately 200 *trans*-acting factors and the incorporation of 79 ribosomal proteins (RPs). The delivery of RPs to pre-ribosomes is a major challenge for the cell because RPs are often highly basic and contain intrinsically disordered regions prone to nonspecific interactions and aggregation. To counteract this, eukaryotes developed dedicated chaperones for certain RPs that promote their solubility and expression, often by binding eukaryotic-specific extensions of the RPs. Rps2 (uS5) is a universally conserved RP that assembles into nuclear pre-40S subunits. However, a chaperone for Rps2 had not been identified. Our lab previously characterized Tsr4 as a 40S biogenesis factor of unknown function. Here, we report that Tsr4 co-translationally associates with Rps2. Rps2 harbors a eukaryotic-specific N-terminal extension that was critical for its interaction with Tsr4. Moreover, Tsr4 perturbation resulted in decreased Rps2 levels and phenocopied Rps2 depletion. Despite Rps2 joining nuclear pre-40S particles, Tsr4 appeared to be restricted to the cytoplasmic. Thus, we conclude that Tsr4 is a cytoplasmic chaperone dedicated to Rps2.

## Introduction

Ribosomes are the complex molecular machines responsible for translation. In eukaryotes, ribosome assembly begins in the nucleolus with the co-transcriptional binding of factors to the pre-rRNA. In total, more than 200 *trans*-acting factors are needed to properly fold, modify, and process the ribosomal RNAs (rRNAs) and to assemble and export the pre-ribosomal subunits to the cytoplasm, where they undergo final maturation (reviewed in (1–4)). Ribosome assembly involves the incorporation of 79 ribosomal proteins (RPs), most of which join the assembling pre-ribosomal particles in the nucleus (5–7). Many ribosomal proteins are poorly soluble as free proteins as they contain disordered regions that adopt their final structure only after delivery to the ribosome. In addition, most ribosomal proteins are RNA binding proteins that are highly positively charged and prone to non-specific interaction with RNA (8). Considering that eukaryotic RPs are synthesized in the cytoplasm and must be transported to the nucleus, their transport to the site of ribosome assembly is particularly challenging given the issues of solubility and non-specific RNA interaction. To deal with these issues cells have evolved dedicated RP chaperones and importins (reviewed in (3, 9)).

Only a limited number of RP chaperones have been identified in yeast (3, 9, 10). Some chaperones are essential (10–14), while the deletion of others give rise to significant growth defects (15–18). These chaperones appear dedicated to their client RPs, and owing to their unique interactions, there is no unified mechanism of binding to their client proteins. For example, Sqt1 uses its WD-repeat β-propeller domain to bind Rpl10 (uL16) (14), Yar1 uses its Ankyrin repeats to bind Rps3 (uS3) (19), and Acl4 uses its TPR repeats to bind Rpl4 (uL4) (20). Reflecting these differences, there is not a universal binding site on the client RPs. Sqt1 and Yar1 recognize the N-termini of their clients (14, 19, 21) while Acl4 binds an internal loop of Rpl4 (15, 16, 20). Despite the diversity of these interactions, all chaperones have a shared role in facilitating RP expression. This is generally achieved via co-translational recognition of the RP by the chaperone (14, 15) to promote RP solubilization by preventing aggregation (10, 14, 15, 17) or to protect RPs from proteasomal degradation (20).

Most RPs are incorporated into pre-ribosomal particles during their assembly in the nucleus. Their nuclear import is dependent on karyopherins (Kaps), also referred to as importins, with the primary RP importins being Kap123 and Kap121 (Pse1) (22), while Kap104 and the Kap95/Kap60 pair have also been implicated in the transport of certain RPs (16, 18, 20, 23). RPs are often bound directly by the importin (10, 16, 20, 22, 24), but it is also possible for a chaperone to act as an import adaptor by facilitating the interaction with the importin. This is the case for Syo1 which simultaneously chaperones Rpl5 (uL18) and Rpl11 (uL5) and serves as an adapter for Kap104 (18, 25). Following nuclear import, the RP-chaperone complex is typically released from the importin in a RanGTP-dependent manner (reviewed in (26)), and the RPs are subsequently loaded onto pre-ribosomes.

Rps2 (uS5) binds to the central pseudoknot (CPK) of the small subunit (SSU or 40S), a critical rRNA feature that establishes the environment for the decoding center (27). Rps2 is needed for the nuclear export of pre-40S particles and their subsequent cytoplasmic maturation involving cleavage of the 20S rRNA intermediate in both budding and fission yeast (6, 28). Despite its role in SSU biogenesis, little is known regarding how Rps2 is chaperoned, imported and delivered to the pre-40S subunit. In humans, an extraribosomal heterotrimeric subcomplex composed of Rps2, the arginine methyltransferase PRMT3, and PDCD2L is needed for pre-40S biogenesis and export, while a paralogous complex of Rps2, PRMT3, and PDCD2 may serve a redundant function (29). Additionally, an orthologous Rps2-PDCD2/Zfrp8 complex in *Drosophila* has been proposed (30). While a chaperone-like function for PDCD2L and PDCD2/Zfrp8 has been suggested (31), it has not been experimentally explored.

We previously identified *TSR4* as an essential 46 kDa protein needed for SSU production (32). Tsr4 shows high sequence similarity to PDCD2L and PDCD2/Zfrp8 (31) suggesting that it may function similarly to these metazoan proteins. Transcriptional repression of *TSR4* results in accumulation of the 20S rRNA processing intermediate, but despite its impact on SSU biogenesis, Tsr4 does not co-sediment with pre-ribosomal complexes (32) hinting that it has an indirect role in 40S maturation. Here, we report that Tsr4 is a dedicated chaperone which co-translationally associates with Rps2 to facilitate its expression. However, unlike its metazoan counterparts, Tsr4 does not appear to enter the nucleus, suggesting that its interaction with Rps2 is restricted to the cytoplasm.

## Results

### N-terminal fragments of Rps2 sequester Tsr4

Structural alignment of Rps2 (uS5) from *S. cerevisiae* 80S (PDB: 4V88) to S5 from *Escherichia coli* 70S (PDB: 4YBB) identified residues 74-231 of yeast Rps2 as constituting the evolutionarily conserved core of the protein, whereas residues 1-73 and 232-254 of Rps2 comprise N- and C-terminal extensions that are not present in the bacterial protein (Fig. 1A). Multiple sequence alignment of Rps2 from various eukaryotic and prokaryotic species revealed that the N-terminal extension is conserved throughout eukaryotic Rps2 proteins (Fig. S1). Although significantly shorter, the C-terminal extension adopts a kinked alpha helix (Fig. 1A) that is conserved among eukaryotic Rps2 proteins (Fig. S1). To explore the functions of the eukaryotic extensions of Rps2, we generated a panel of seven GFP-tagged Rps2 truncations expressed under the control of its native promoter on centromeric vectors (Fig. 1A). Expression of Rps2_(1-73)_ or Rps2_1-223_, containing residues 1-73 and 1-223, respectively, was strongly dominant negative (Fig. 1B) despite their expression from low copy vectors using the native *RPS2* promoter. Expression of the slightly longer Rps2_1-235_ fragment was only slightly dominant negative, and this was the only truncation mutant that was able to complement loss of *RPS2*, to any degree (Fig. S2A). Interestingly, the dominant negativity of Rps2_1-73_ was partially alleviated by the addition of the C-terminal extension (aa 224 to 254; Rps2_Δ74-223_) (Fig. 1B). Expression of the other fragments of Rps2 did not cause any obvious growth defect, and western blotting showed that the dominant negative effect of expressing the N-terminal fragments did not correlate with higher protein expression (Fig. S2B). Thus, the dominant negative effect of the N-terminal fragments is most easily explained by sequestering a factor or set of factors.

**Figure 1.**
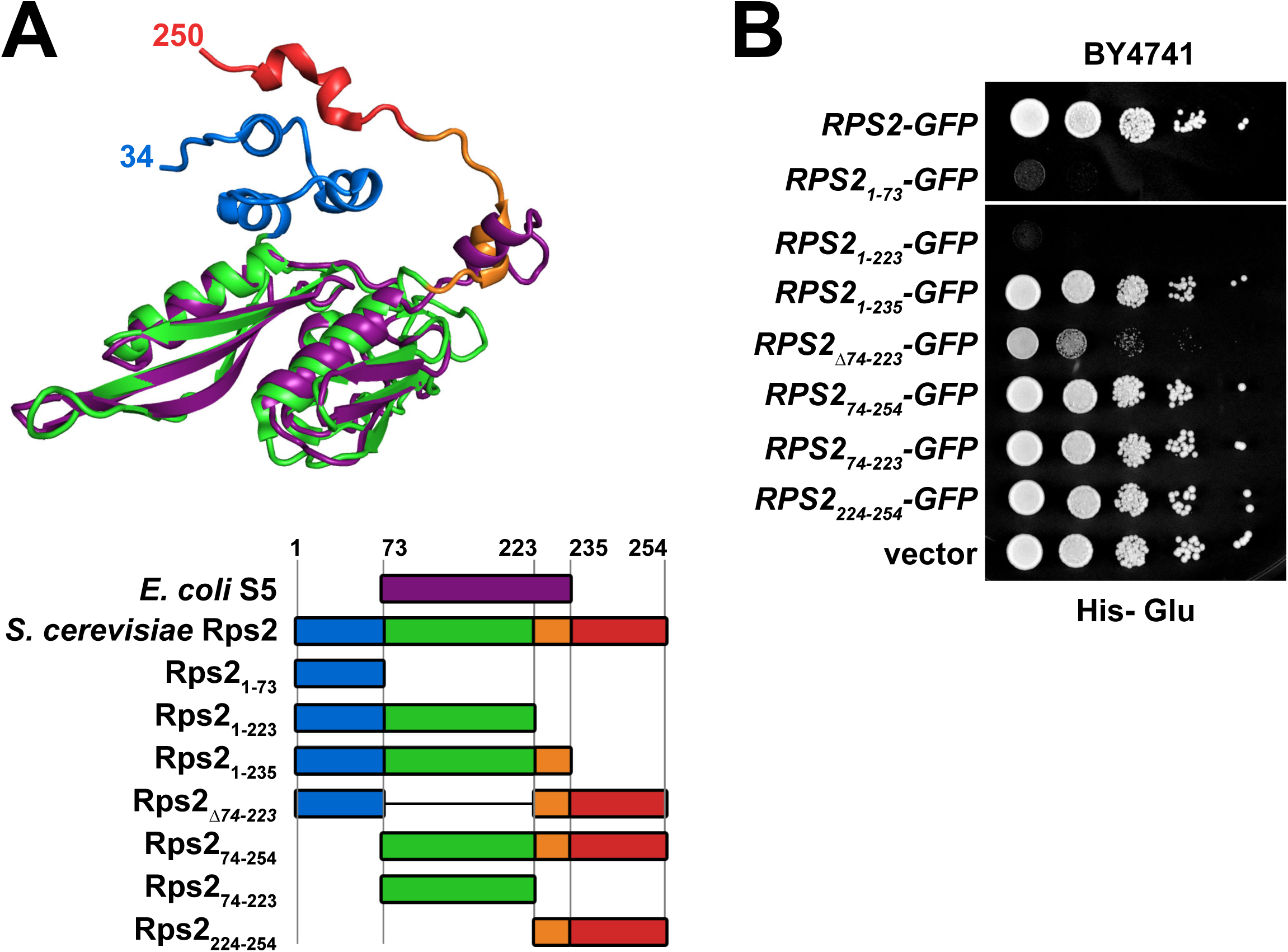
C-terminal truncations of Rps2-GFP are strongly dominant negative. A) Top: Structural alignment of *E. coli* S5 (purple, from PDB 4YBB) with yeast Rps2 (uS5) (multicolored, from PDB 4V88) generated with PyMol. Bottom: cartoon depiction of the alignment of full-length Rps2 and deletion mutants with *E. coli* S5, colored as for the structural alignment. Amino acid positions are relative to yeast Rps2. B) Wild-type *RPS2* and the indicated deletion mutants were ectopically expressed as GFP fusions in wild-type (BY4741) cells under control of the *RPS2* promoter. 10-fold serial dilutions were plated on glucose-containing media lacking histidine and imaged after 2 days at 30°C.

To identify potential binding partners of the fragments of Rps2-GFP, we expressed each fragment under the control of the galactose-inducible *GAL1* promotor in a *reg1-501* mutant strain that allows galactose induction in the presence of glucose (33). After brief expression of the Rps2-GFP fragments, extracts were prepared, and ribosomes were removed by ultracentrifugation prior to immunoprecipitation (IP) of the Rps2-GFP fragments to identify extraribosomal binding partners. Each fragment showed a slightly different set of co-purifying binding partners (Fig. 2A). However, some co-purifying species were common to the N-terminal fragments that were not present in the C-terminal fragments and *vice versa*. The N-terminal fragments all co-immunoprecipitated a species that migrated slightly above 50 kDa on SDS-PAGE (Fig. 2A; lanes 2-4) and was identified as Tsr4 by mass spectrometry. Although full-length Rps2-GFP co-migrated with Tsr4 and masked its signal in SDS-PAGE (Fig. 2A, lane1), Tsr4 was detected by mass spectrometry in the wild-type sample as well. To independently verify that the N-terminus of Rps2 facilitates its interaction with Tsr4, we performed the converse immunoprecipitation experiment. To this end, we genomically tagged *TSR4* with *TEV-13myc* in the *reg1-501* strain and briefly expressed the various Rps2-GFP fragments. We also expressed GFP alone as a negative control. Extracts were prepared and immunoprecipitations were done using Tsr4-TEV-13myc as bait, and western blotting was used to detect co-precipitation of the GFP-tagged Rps2 fragments or GFP alone. The Rps2-GFP fragments harboring the N-terminus co-precipitated with Tsr4 more abundantly than a fragment lacking the N-terminus, while GFP alone did not co-precipitate, indicating specificity for the Rps2 fragments (Fig. 2B). This result suggests that while Tsr4 has some affinity for the core of Rps2, the N-terminus is its primary binding site.

**Figure 2.**
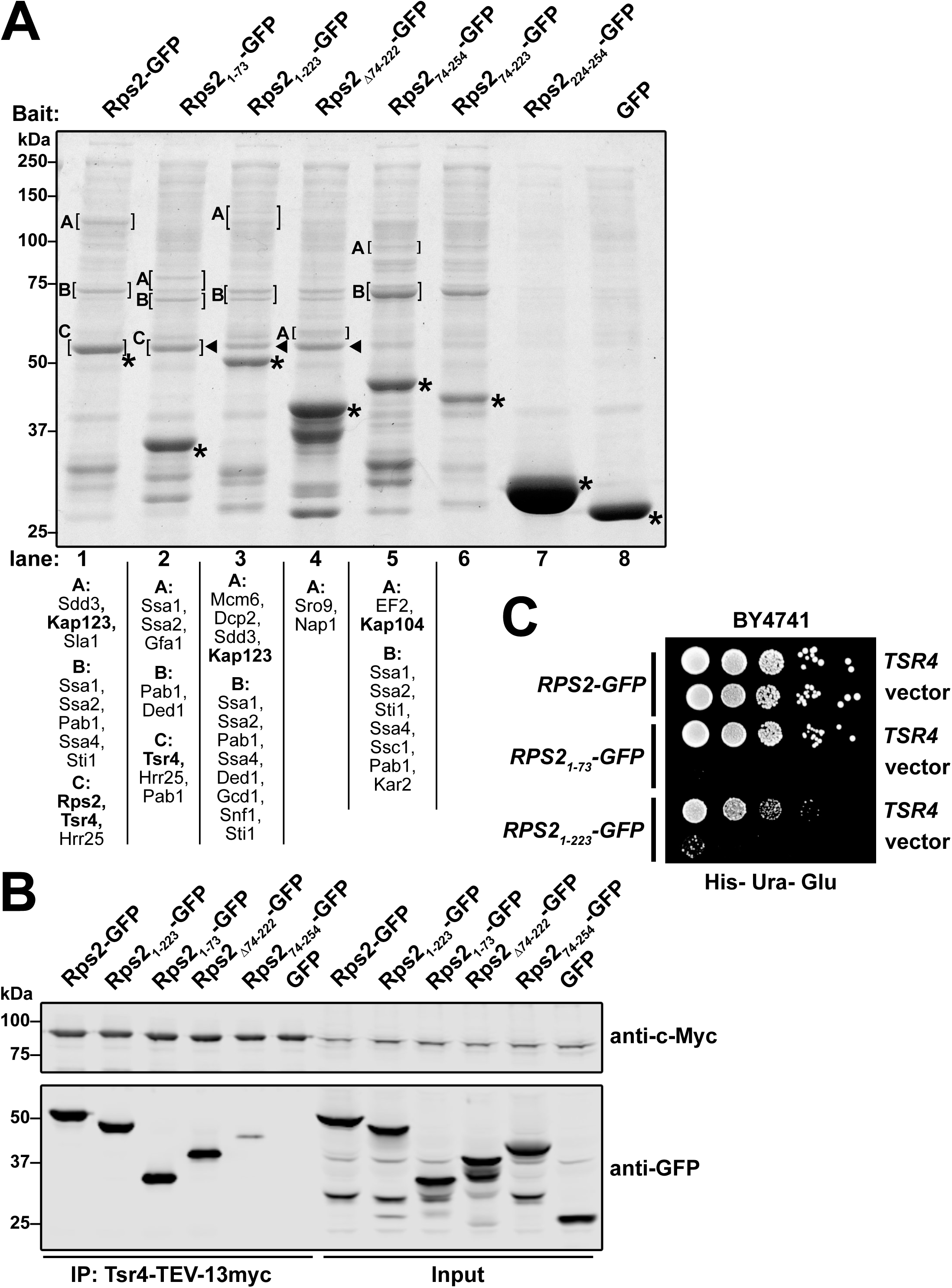
C-terminal truncations of Rps2-GFP sequester Tsr4. A) The indicated Rps2-GFP proteins and GFP alone were expressed in strain AJY1134. Extracts were cleared of ribosomes by ultracentrifugation. GFP fusion proteins were immunoprecipitated and analyzed by SDS-PAGE and staining with Coomassie blue. The indicated bands (brackets) were excised for mass spectrometry. Proteins identified with greater than 50 total peptides are listed below each lane ordered by abundance. Proteins of particular interest are indicated by bold text. Asterisks (*): bait proteins; triangles (◂): Tsr4. B) The indicated Rps2-GFP fragments and GFP alone were expressed in AJY4363. Extracts were generated and Tsr4-TEV-13myc was immunoprecipitated. The co-precipitating proteins were separated by SDS-PAGE and analyzed by Western blotting probing with anti-*c*-Myc and anti-GFP. C) The toxic C-terminal truncations of Rps2-GFP, Rps2_1-73_ and Rps2_1-223_, were co-expressed in wild-type (BY4741) cells with high-copy *TSR4* or empty vector. 10-fold serial dilutions were plated on glucose-containing media lacking histidine and uracil and grown for 3 days at 30°C.

*TSR4* is an essential gene that was first identified in a bioinformatic analysis of novel ribosome biogenesis factors and is required for the processing of the 20S rRNA intermediate (32). However, its specific function was not further explored. Because the Rps2 fragments harboring the N-terminus all associated with Tsr4, while the fragments lacking it did not, we speculated that the N-terminal fragments were dominant negative due to sequestration of Tsr4. Consistent with this notion, the dominant negative defect of Rps2_1-73_ was fully alleviated by expression of *TSR4* from a high-copy vector (Fig. 2C). Surprisingly, the dominant negative effect of the longer Rps2_1-223_ fragment was only partially suppressed by overexpression of *TSR4* suggesting that it has a negative effect in addition to Tsr4 sequestration. Quantitative Western blots using a GFP-tagged version of the 60S biogenesis factor Nmd3 for a standard curve indicated that the amount of Tsr4-GFP is roughly one-fifth the amount of Nmd3-GFP (data not shown). On average, there are approximately 7,500 Nmd3 molecules per cell (34). Thus, we estimate that each cell has about 1,500 Tsr4 molecules. This number is low compared to other RP chaperones like Yar1 and Sqt1 which are estimated to be present at about 11,000 and 7,600 molecules per cell, respectively (34). Consequently, low expression of Rps2_1-73_ or Rps2_1-223_ could effectively sequester the limited amount of Tsr4 available in a cell and have a profound effect on cell growth.

Because the Rps2-GFP immunoprecipitations (Fig. 2A) were done on samples pre-cleared of ribosomes, we next asked if the most toxic N-terminal fragments of Rps2 could associate with ribosomes. Full-length and N-terminal fragments of Rps2 were expressed from centromeric vectors under the control of their cognate promoter in a *P*_*GAL1*_*-RPS2* strain in which endogenous *RPS2* expression was repressed with glucose for 5 hours to eliminate competition between endogenous *RPS2* and the plasmid-borne constructs. Extracts were subjected to ultracentrifugation through sucrose density gradients to separate extraribosomal proteins from ribosome-bound proteins. Full-length Rps2-GFP co-sedimented with 40S, 80S, and polysomes with very little present as free protein (Fig. 3A) consistent with it being a functional protein (Fig. S2A) that is incorporated into ribosomes. In contrast, Rps2_1-73_ was present exclusively at the top of the gradient, indicating that it does not stably associate with ribosomes (Fig. 3B). A portion of the Rps2_1-223_ pool was also present at the top of the gradient but, unlike Rps2_1-73_, the larger Rps2_1-223_ fragment also co-sedimented with 40S subunits and 80S ribosomes in sucrose density gradients with very small quantities detected in polysomes (Fig. 3C). It seems likely that the incorporation of the truncated Rps2_1-223_ fragment into ribosomes disrupts either their maturation or function, accounting for the observation that the dominant negative effect of this fragment cannot be fully suppressed by overexpression of TSR4 (Fig. 2B). Notably, the UV-trace from the cells expressing Rps2_1-223_-GFP showed a deficit of 40S relative to 60S subunits compared to WT Rps2-GFP, indicating a defect in 40S biogenesis (compare Fig. 3C to 3A, black trace). The ability of N-terminal fragments of Rps2 to titrate out Tsr4, blocking its function in ribosome assembly and leading to a dominant negative phenotype, is reminiscent of genetic interactions between some RPs and their chaperones (14, 15, 21) and led us to hypothesize that Tsr4 is a dedicated chaperone for Rps2.

**Figure 3.**
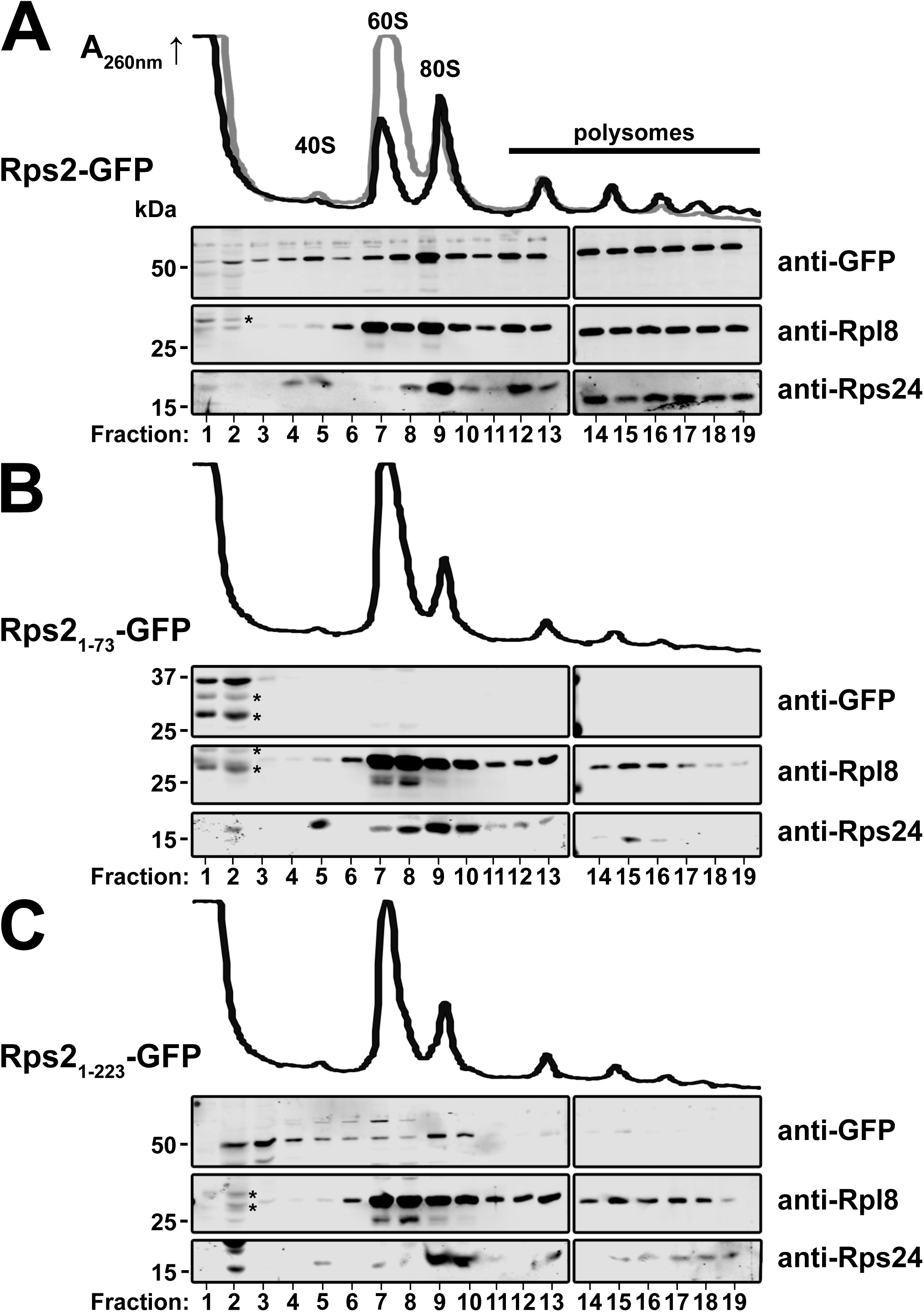
Sucrose density gradient sedimentation of Rps2-GFP fragments. The *P*_*GAL1*_*-RPS2* strain AJY3398 containing empty vector or vectors expressing GFP fusions of the indicated C-terminal Rps2 truncations under control of the cognate promoter was cultured in glucose containing medium for 5 hours to repress genomic *RPS2* expression prior to a 10 min treatment with cycloheximide. Extracts were generated and fractionated on sucrose density gradients and proteins were precipitated. The sedimentation of relevant proteins was monitored by western blotting following SDS-PAGE. Traces of continuous UV absorbance at 260 nm are shown. A) Cells expressing wild-type Rps2-GFP (black) trace or no Rps2 (gray). B and C) cells expressing Rps2_1-73_-GFP and Rps2_1-223_-GFP, respectively. Degradation products of the Rps2-GFP fragments are denoted by asterisks (*).

### Residues within the extreme N-terminus and a helical bundle of Rps2 facilitate the interaction with Tsr4

Because the genetic interaction between *RPS2* and *TSR4* likely reflects a physical interaction, we sought to more finely map this interaction by identifying residues of Rps2 that are necessary for its interaction with Tsr4. We reasoned that we could use the dominant negative growth phenotype of Rps2_1-73_ as a proxy for its interaction with Tsr4. Thus, *Rps2*_*1-73*_ mutants that alleviate its toxicity would likely disrupt its interaction with Tsr4. To this end, we used error-prone PCR to generate a pool of amplicons containing random mutations in *RPS2*_*1-73*_*-GFP*. We recombined the amplicons into a *GAL1*-inducible expression vector *in vivo* by homologous recombination. We plated the library on galactose-containing media and screened for relief of the dominant negative growth phenotype. We isolated 110 colonies from approximately 3,000 transformants with varying degrees of improved growth, of which 73 showed GFP fluorescence indicating expression of the fusion protein. Sanger sequencing identified 48 single point mutations within the *RPS2*_*1-73*_ coding region revealing 35 unique mutations that alleviated the dominant negativity of *RPS2*_*1-73*_ (Fig. 4A and 4B and data not shown).

**Figure 4.**
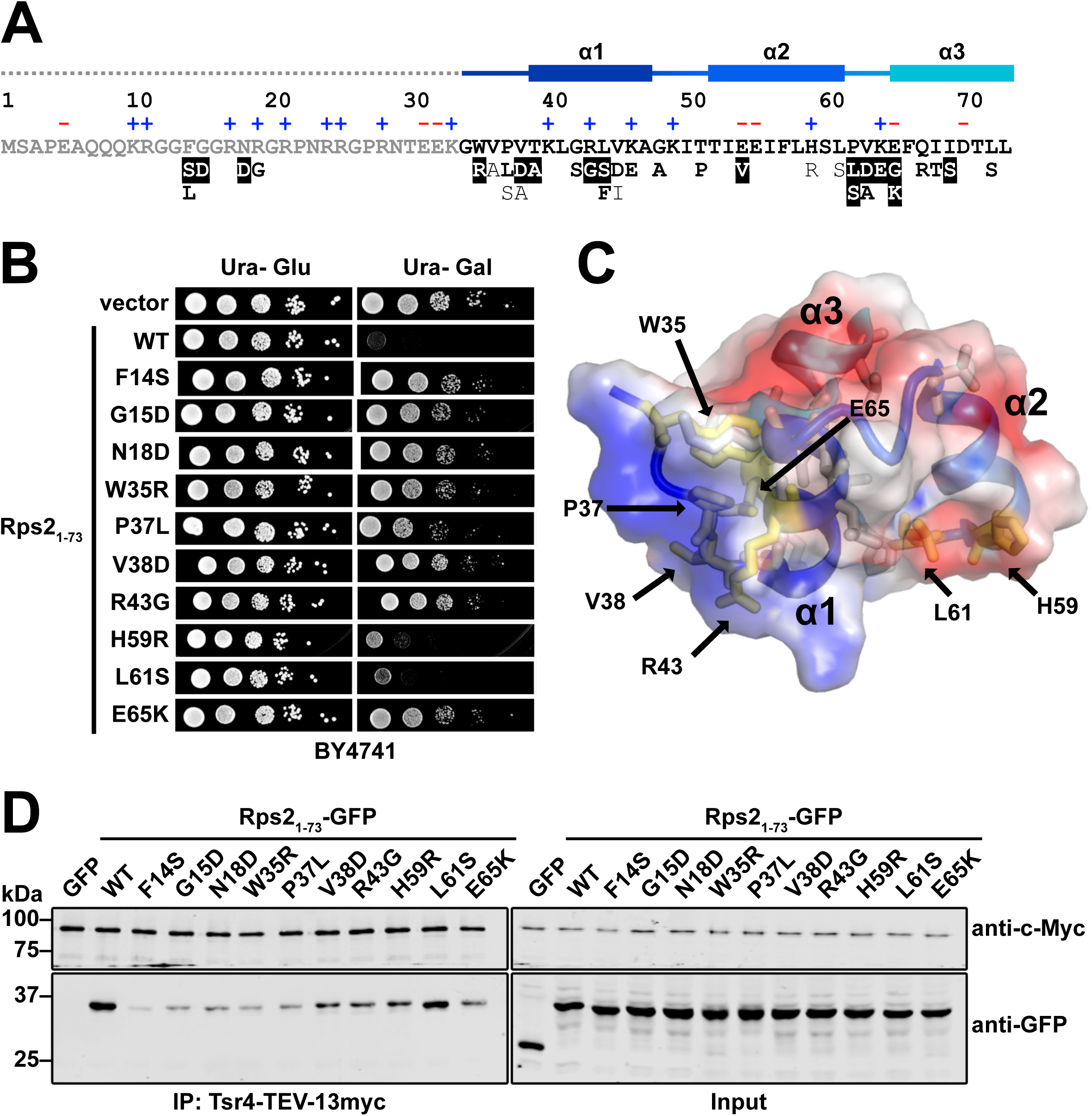
Tsr4 recognizes two regions of the N-terminus of Rps2. A) Primary sequence of the N-terminal 73 residues of yeast Rps2 with charged residues indicated (+,-) and secondary structure elements from PDB 4V88 indicated above. Gray dashed line not resolved in PDB 4V88. Mutations that alleviated the dominant negative growth phenotype of Rps2_1-73_-GFP are listed below the primary sequence. Strong mutants (white text with black highlight), moderate mutants (bold, black text), and weak mutants (black text) are indicated below. B) Growth of a subset of mutants that relieved the dominant negative phenotype of Rps2_1-73_-GFP. Ten-fold serial dilutions of wild-type (BY4741) cells expressing the indicated mutants under control of the galactose-inducible *GAL1* were plated on glucose- or galactose-containing media lacking uracil and grown for 2 days at 30°C. C) The helical bundle of Rps2 residues 34-73 is shown in cartoon with transparent electrostatic surface (from PDB: 4V88). Amino acids that can be mutated to alleviate the growth inhibition of Rps2_1-73_ are shown as sticks with residues included in panel B shown in yellow. D) Strength of interaction between Tsr4 and wild-type (WT) Rps2_1-73_-GFP or the indicated Rps2_1-73_-GFP mutants as determined by their ability to co-immunoprecipitate with Tsr4-TEV-13myc.

Residues 1-33 of the N-terminal extension of Rps2 contain an RG-rich motif (Fig 3A) commonly found in intrinsically disordered regions of RNA-binding proteins (35). Consistent with its predicted disordered nature, this region of Rps2 has not been resolved in crystal or cryo-EM structures of 40S subunits. Five of the unique mutations identified in our screen resulted in substitutions within the RG-rich motif (Fig. 4A). Residues 34-73 fold into three short alpha helices that form a small helical bundle (Figs. 4A and 4C). The remaining 30 mutated residues were within this helical bundle (Figs. 4A and 4C) and the majority of these mapped to a positively charged surface (Fig 4C). These results suggest that Tsr4, which has an overall net negative charge (pI = 4.5), binds the N-terminal extension of Rps2, in part through electrostatic interactions.

To determine if these mutants lose interaction with Tsr4, we chose ten representative mutants to investigate further (Fig. 4B). Of these, seven mapped to the N-terminal helical bundle of Rps2 (Fig 4C; yellow residues). We expressed wild-type and mutant *RPS2*_*1-73*_*-GFP* under control of the galactose-inducible *GAL1* promoter in the *reg1-501* strain containing genomic *TSR4-TEV-13myc*. As a negative control, we expressed GFP alone. After a brief galactose induction, extracts were prepared, Tsr4-TEV-13myc was immunoprecipitated and western blotting was used to detect co-precipitation of the Rps2_1-73_-GFP proteins or GFP. Wild-type Rps2_1-73_-GFP showed specific association with Tsr4 as GFP alone was not detected in the Tsr4 immunoprecipitation (Fig. 4D, left panel). All of the mutants showed reduced association with Tsr4 compared to wild-type Rps2_1-73_. The apparent differences in binding were not due to variable expression of the mutant proteins, as all showed similar levels of expression (Fig. 4D, right panel). In general, the degree of relief of the dominant negative phenotype correlated with loss of Tsr4-Rps21-73 interaction. In addition, the N-terminal-proximal mutations appeared to have a stronger impact on Tsr4 interaction than do mutations further from the N-terminus.

### Tsr4 co-translationally associates with Rps2 and is needed for its efficient expression

Dedicated RP chaperones often bind their client proteins co-translationally, soon after their interaction domain is presented from the exit tunnel of the ribosome (14, 15). For example, Sqt1 is a chaperone for Rpl10 (uL16) that co-translationally associates with the N-terminus of Rpl10 to allow copurification of *RPL10* mRNA (14). Based on our observations that Tsr4 interacts with the N-terminus of Rps2, we speculated that Tsr4 could similarly bind Rps2 co-translationally. In order to test this idea, we affinity purified *Tsr4-TAP* and *Sqt1-TAP* and probed for specific mRNAs using RT-qPCR. Cells were treated with cycloheximide (CHX) to arrest translation elongation. Extracts were made, and affinity purifications were done in the presence of CHX. We isolated the co-purifying RNA and used it as template in RT-qPCR reactions, amplifying either *RPS2* or *RPL10* mRNA for both the Tsr4 and Sqt1 purifications. We expected that Tsr4 would efficiently pull down *RPS2* mRNA but not *RPL10* mRNA, while Sqt1 would specifically purify *RPL10* mRNA but not *RPS2* mRNA. To display enrichment relative to the expected target mRNA of each bait, we first normalized the number of mRNA molecules isolated from immunoprecipitations, to the number of mRNA molecules in the input, then set this value for the expected target mRNA for each bait to 1.0 (Fig. 5). This analysis revealed that Tsr4-TAP enriched specifically for *RPS2* mRNA but not *RPL10* mRNA. Conversely, Sqt1-TAP enriched for *RPL10* mRNA, but not *RPS2* mRNA. We also performed the experiment using Tsr4-GFP as bait and again saw an enrichment of the *RPS2* mRNA over the *RPL10* mRNA. Thus, we conclude that Tsr4 associates with Rps2 co-translationally.

**Figure 5.**
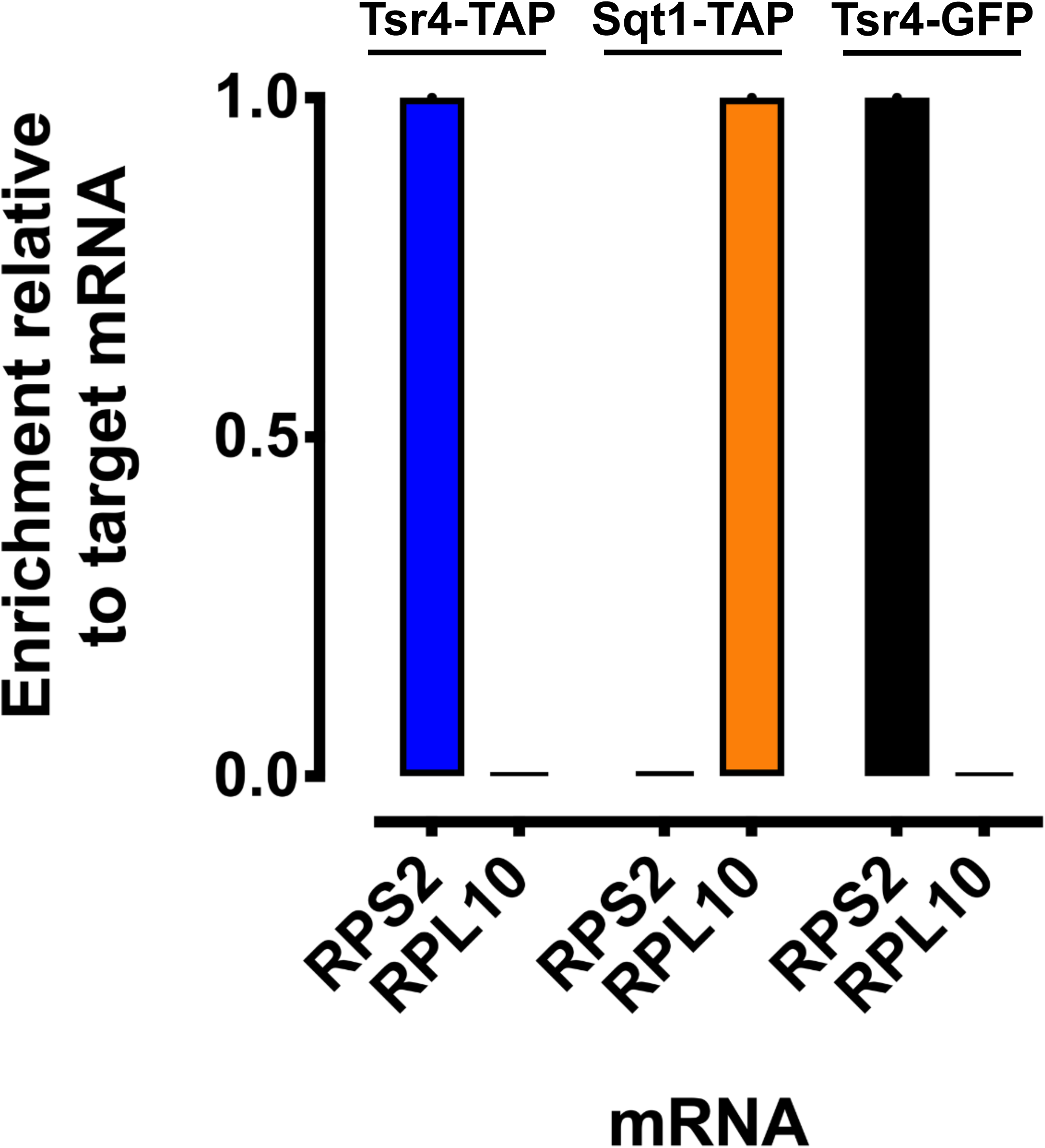
Tsr4 associates co-translationally with nascent Rps2. The co-translational association of Tsr4 and Sqt1 with their client proteins, Rps2 and Rpl10, respectively, was determined by their ability to co-immunoprecipitate the mRNA of their client protein as measured by RT-qPCR. To prevent ribosome run-off of mRNAs, cells were treated with CHX prior to harvest, and extracts and immunoprecipitations were done in the presence of CHX. Abundance is shown as mRNA enrichment relative to the expected target mRNA of each bait. The qPCR reactions were done in technical triplicates; error bars indicate the standard deviation for the ratios of mRNA molecules in the IP to Input for each replicate. See “Materials and Methods” for additional details.

Chaperones often facilitate the expression of their target RPs (10, 13, 14, 17). To test if Tsr4 similarly enhances the expression of Rps2, we transformed a vector encoding Rps2-GFP under the control of a galactose-inducible promoter into strains harboring either a wild-type or temperature-sensitive allele of Tsr4 (*tsr4-ts*). The *tsr4-ts* mutant was unable to grow at the non-permissive temperature of 37°C but was viable at 30°C and room temperature (∼25°C). Cells were cultured in raffinose-containing media at room temperature and then shifted to 37°C for 2 hours to inactivate the temperature-sensitive Tsr4 protein. Expression of *Rps2-GFP* was then briefly induced by the addition of galactose for 90 minutes. Extracts were prepared and free proteins were separated from ribosomes by ultracentrifugation. Western blotting was performed on the input, supernatant, and the pellet. This analysis showed that the levels of Rps2-GFP in the input, supernatant, and pellets were reduced in the strain background harboring *tsr4-ts* relative to the strain harboring the wild-type *TSR4* allele (Fig. 6A). This decrease of Rps2-GFP was also apparent when cells were imaged by fluorescent microscopy after 60 minutes of induction (Fig. 6B). Together, these results suggest that functional Tsr4 is needed for the efficient expression of Rps2.

**Figure 6.**
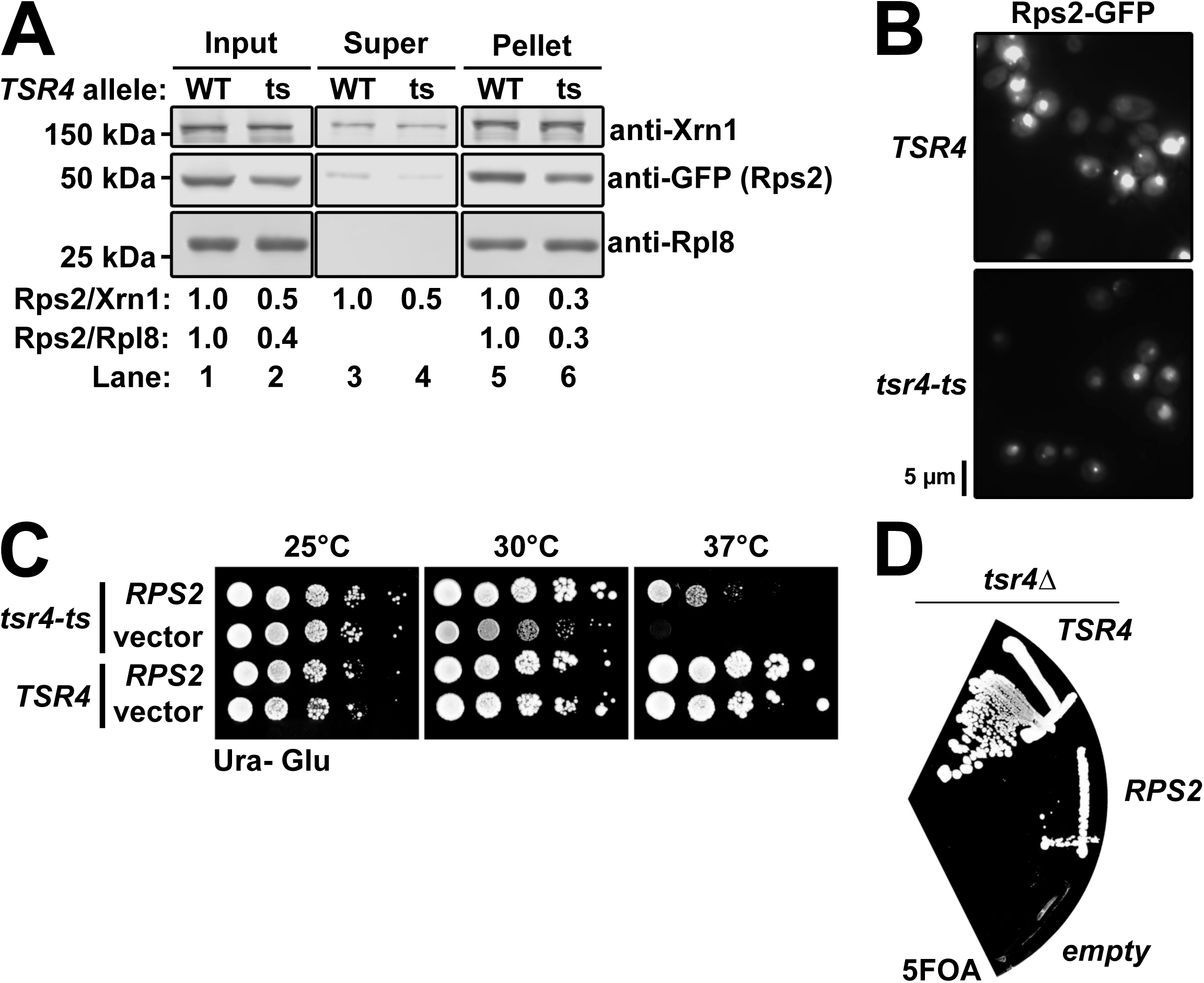
Tsr4 is needed for efficient expression of Rps2. A) The expression of Rps2-GFP was analyzed in *tsr4-ts* (AJY2873) cells compared to wild-type (WT) *TSR4* cells (BY4741). Cells were shifted to non-permissive temperature for 2 hours prior to the expression of Rps2-GFP for 90 minutes by the addition of galactose. Comparable amounts of extract were overlaid onto sucrose cushions prior to ultracentrifugation to separate free proteins (supernatant) from ribosomes (pellet). Rpl8 and Xrn1 were used as loading controls. The relative levels of Rps2 are quantified below, as the ratio of Rps2 to Xrn1 and the ratio of Rps2 to Rpl8, where the ratio for the wild-type sample was set to one. B) GFP fluorescent microscopy of *TSR4* or *tsr4-ts* cells expressing Rps2-GFP. Cells were cultured as described for panel A, except Rps2-GFP was induced for 60 minutes after the 2-hour shift to 37°C. C) Ten-fold serial dilutions of *TSR4* (BY4741) and *tsr4-ts* (AJY2873) cells transformed with either empty vector (pRS416) or a vector encoding *RPS2* (pAJ4244) under the transcriptional control of its cognate promoter after 3 days at the indicated temperatures on SD-ura media containing glucose. D) *tsr4*Δ cells (AJY4359) containing a *URA3*-marked *TSR4* vector were transformed with *HIS3*-marked vectors encoding *TSR4* (pAJ4232) or *RPS2* (pAJ4207) or an empty vector (pRS416) and complementation was assessed by plasmid shuffle on 5FOA-containing media after 6 days of growth.

The need for a dedicated chaperone may be suppressed by increased production of its client protein (10, 14–17). We asked if the *tsr4-ts* mutant could be suppressed by increased gene dosage of *RPS2*. Indeed, excess *RPS2* expressed from a centromeric vector fully suppressed the slow grow defect of *tsr4-ts* at 30°C and partially suppressed its lethality at 37°C (Fig. 6C). This suggests that increased expression of Rps2 stabilizes the product of *tsr4-ts* and/or bypasses the need for Tsr4 altogether. To determine if overexpression of *RPS2* could bypass the lethality of a *tsr4*Δ mutant, we used a plasmid shuffle technique. We introduced *HIS3*-marked plasmids encoding *TSR4, RPS2*, or nothing (empty vector) into a *tsr4*Δ strain harboring a *URA3*-marked vector encoding *TSR4*. We then assayed for the ability of these *HIS3*-marked vectors to shuffle out the *URA3*-marked vector by their growth on media containing the 5-Fluoroorotic acid (5FOA). As expected, the *HIS3*-marked *TSR4* vector complemented *tsr4*Δ (Fig. 6D). The *RPS2* vector weakly suppressed the lethality of *tsr4*Δ, while the empty vector did not support growth. We confirmed the absence of *TSR4* in the *RPS2*-encoding cells by PCR (data not shown). Thus, increased gene dosage of *RPS2,* and a presumably concomitant increase in production of Rps2 protein, can bypass the essential function of Tsr4, consistent with the notion that Tsr4 is a chaperone for Rps2.

### Repression of *TSR4* expression blocks pre-40S export

Pre-40S particles exported from the nucleus contain 20S pre-RNA. In the cytoplasm, the endonuclease Nob1 cleaves 20S rRNA into the mature 18S (36, 37), liberating a small 5’ fragment of the internal transcribed spacer 1 (ITS1) that is rapidly degraded by the exoribonuclease Xrn1 (38). Previous studies in *S. cerevisiae and S. pombe*, using fluorescent *in situ* hybridization (FISH) with an oligo probe specific to the D-A2 region of ITS1 of rRNA intermediates or using fluorescent microscopy using Rps7-GFP as a marker for 40S subunits, showed that depletion of Rps2 prevents pre-40S export (6, 28). If Tsr4 is a chaperone for Rps2 then one would expect that the depletion of Tsr4 should phenotypically mimic depletion of Rps2. However, our lab previously reported that depletion of Tsr4 does not block the export of pre-40S particles from the nucleus (32), but that study used Rps2-GFP as a marker for pre-40S. Because Tsr4 is needed for the expression of Rps2 (Fig. 6A), the depletion of Tsr4 leads to loss of newly synthesized Rps2-GFP, rendering Rps2-GFP ineffective as a reporter for the role of Tsr4 in pre-40S export.

To revisit the question of whether or not Tsr4 is important for pre-40S export, we generated strains in which *RPS2* or *TSR4* were placed under the transcriptional control of the glucose-repressible *GAL1* promoter in an *xrn1*Δ background. Deletion of *XRN1* allows the liberated ITS1 fragment to accumulate in the cytoplasm, providing a clearer change in FISH signal when export is blocked (39). The transcription of *RPS2* or *TSR4* was repressed for 2 hours by the addition of glucose after which cells were prepared for FISH using an oligo probe that hybridizes specifically to the ITS1 region. As expected, signal for ITS1 was predominantly cytoplasmic in the *xrn1*Δ strain that expresses both *RPS2* and *TSR4* (Fig. 7). Upon repression of *RPS2* or *TSR4* transcription, the signal for ITS1 accumulated in the nucleoplasm, indicating a defect in pre-40S export. Thus, the transcriptional repression of Tsr4 phenocopies the transcriptional repression of Rps2 and Tsr4 is needed for pre-40S export.

**Figure 7.**
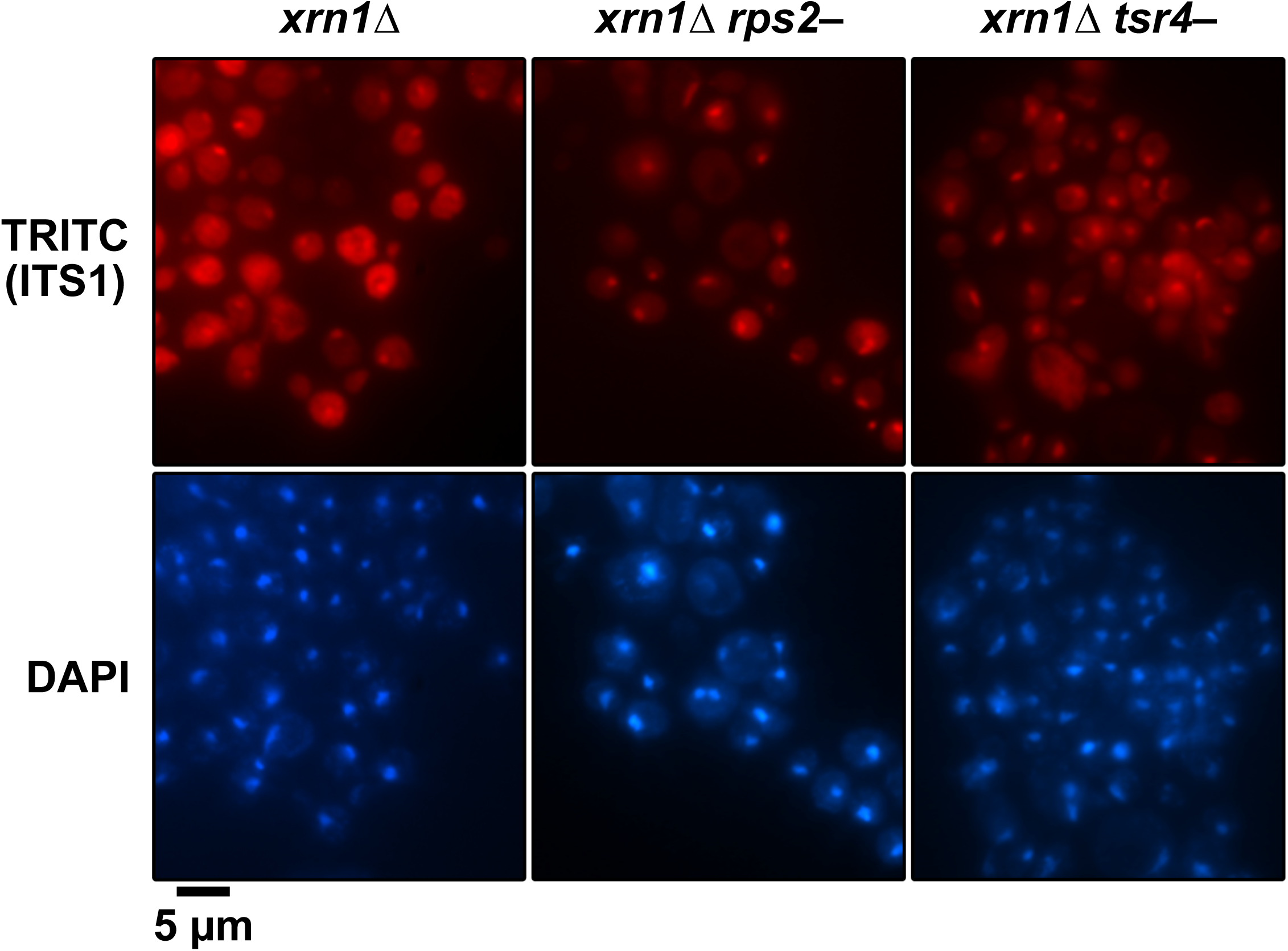
Repression of *TSR4* blocks the nuclear export of pre-40S. The subcellular distribution of ITS1 was analyzed by FISH with a Cy3-labeled oligonucleotide complementary to the D-A2 region of ITS1 (TRITC) in *xrn1*Δ (RKY1977), *P*_*GAL1*_*-RPS2 xrn1*Δ (AJY4300), and *P*_*GAL1*_*-3xHA-TSR4 xrn1*Δ (AJY4297) cells after switch to glucose-containing media for two hours. The position of the nucleus was determined by DAPI staining and overall cell outline by differential interference contrast (DIC).

### Tsr4 appears restricted to the cytoplasm

Several dedicated RP chaperones enter the nucleus with their client RPs (13, 15, 17, 18) to prevent their interaction with RNA cargo being exported from the nucleus and to facilitate loading of the RP onto pre-ribosomes once inside the nucleus. Tsr4 contains a putative, canonical bipartite nuclear localization signal (NLS) sequence (Fig. S3A) suggesting that Tsr4 may usher Rps2 into the nucleus. However, mutants containing alanine substitutions of critical lysine residues of the putative NLS (Tsr4_ΔNLS_-GFP) fully complemented the loss of Tsr4 (Fig. S3A), suggesting that this NLS is not functional or not important for function. To further test the possibility that Tsr4 shuttles between the nucleus and the cytoplasm, we blocked Crm1 (Xpo1)-dependent nuclear export. Crm1 is a nuclear export receptor for pre-ribosomal subunits and proteins containing leucine-rich nuclear export sequences (reviewed in (40)). The Crm1_T539C_ variant harbors a mutation that sensitizes it to inhibition by Leptomycin B (LMB) (41). Tsr4-GFP did not accumulate in the nucleus following treatment with LMB suggesting that it does not shuttle between the nucleus and cytoplasm in a Crm1-dependent manner (Fig. S3B). These data lead us to conclude that it is unlikely that Tsr4 enters the nucleus and that Rps2 must be handed off to a karyopherin prior to its nuclear import.

### Core-containing fragments of Rps2 are bound to importins

Kap123 is considered the primary importin for RPs (22) while Kap104 has been shown to interact with specific RPs (16, 18, 20, 24). We identified unique spectra for Kap123 and Kap104 in the co-immunoprecipitation experiments of core-containing Rps2-GFP fragments, but the purifications of the fragments lacking the core lacked any corresponding bands for these proteins (Fig. 2A). This result suggests that recognition of the Rps2 core leads to the recruitment of an importin. Consistent with this idea, the subcellular distributions of the core-containing fragments that lacked either the N-terminus or the C-terminus or lacked both termini were nuclear (Fig. S4), while the Rps2_1-73_-GFP and the Rps2_224-254_-GFP fragments were not. An exception to this is that Rps2_1-235_-GFP was found in the cytoplasm. However, this mutant was slightly functional (Fig. S2A) suggesting that this signal is probably due to it being in translating ribosomes. Contrary to the notion that the core of Rps2 is needed for its import, the Rps2_Δ74-223_-GFP fragment, which lacks the core region, was also nuclear (Fig. S4), suggesting some redundancy for Rps2 import that relies on recognition of the C-terminal portion of Rps2.

## Discussion

Rps2 is a universally conserved RP, but eukaryotic Rps2 contains unique N- and C-terminal extensions. Here, we provide evidence that these eukaryotic extensions of Rps2 drive its interaction with Tsr4, which we identify as a dedicated cytoplasmic chaperone for Rps2. Like other chaperones of RPs, Tsr4 finds its client protein co-translationally, recognizing the N-terminal extension of Rps2 as it emerges from the ribosome. After translation, Tsr4 is released from Rps2 in the cytoplasm by an unknown mechanism (see below). We speculate that either concurrently or soon after Tsr4 release an importin binds to the conserved core of Rps2 and brings it to the nucleus where it is delivered to pre-40S particles.

### How does Tsr4 facilitate Rps2 expression?

Tsr4 is a modestly expressed protein which is easily titrated by even low levels of the N-terminal fragment of Rps2. This raises the question of how the small pool of Tsr4 finds nascent Rps2 in the large pool of translating ribosomes at a rate to sustain the active ribosome assembly pipeline? One possibility is that Tsr4 also has some affinity for the *RPS2* mRNA which helps localize it to the site of Rps2 translation. It is also possible that Tsr4 assembly with the N-terminus of Rps2 itself is promoted by additional factors. The Ccr4-Not complex has recently been implicated in co-translational assembly of the regulatory cap of the proteasome (42). Intriguingly, *TSR4* was identified in a screen that also identified multiple components of the NOT complex, suggesting a functional link between *TSR4* and the Ccr4-Not complex (43). Alternatively, the limiting levels of Tsr4 in cells and rate at which it engages with nascent Rps2 may set the level of Rps2. Excess RPs, including Rps2, are rapidly targeted for proteasomal degradation by the ubiquitin ligase Tom1 and become insoluble in the absence of Tom1 (44, 45). Thus, Rps2 may be constitutively expressed and degraded and only the population that is captured co-translationally by Tsr4 is competent for assembly into pre-40S particles in the nucleus. A similar model for the co-translational assembly of multimeric protein complexes has recently been proposed as a means to protect highly basic or hydrophobic regions of proteins prior to assembly (46).

### How is Rps2 handled after Tsr4 is released?

Our data suggests that Tsr4 does not enter into the nucleus (Fig. S3), indicating that Rps2 is released from Tsr4 and handed off to an import factor. This raises the question of how is Tsr4 released from Rps2? When bound to the ribosome, the N-terminus of Rps2 makes an intramolecular interaction with its C-terminus (Fig. 1A) (27). Because Tsr4 binds to the N-terminus of Rps2, a tempting model is that the C-terminus facilitates release. Interestingly, fusion of the C-terminus to the N-terminal fragment of Rps2 partially alleviated the toxicity of the N-terminal fragment (Fig. 1B; *cf*. Rps2_1-73_ to Rps2_Δ74-223_). However, we have not been able to demonstrate that the C-terminus of Rps2 affects the interaction with Tsr4 in vitro (Fig. 2B and data not shown). Subsequent to or concomitant with the release from Tsr4, Rps2 must bind an importin, likely Kap123 and/or Kap104 to be ferried into the nucleus.

### Differences between S. cerevisiae Tsr4/Rps2 and homologous systems

Tsr4 is evolutionary conserved amongst eukaryotes (31). PDCD2/Zfrp8 in *Drosophila* directly interacts with Rps2 and appears to serve a function in 40S production, but this interaction has not been extensively studied (30). Humans have two Tsr4 homologs, PDCD2 and PDCD2L (31). The Bachand group showed that these two proteins have somewhat redundant roles in 40S production (29), however PDCD2L was characterized more extensively than PDCD2. Human PDCD2L is needed for Rps2 expression, consistent with our results with yeast Tsr4, suggesting that the human protein acts as a chaperone for Rps2. However, unlike yeast Tsr4, PDCD2L shuttles into the nucleus and binds to pre-40S particles (29, 32). PDCD2L also harbors a leucine-rich nuclear export signal, suggesting that it may also function as an export adaptor (29). Interestingly, while PDCD2L binds pre-40S subunits, PDCD2 does not. Moreover, knock-down of PDCD2 has a greater effect on 40S production than knock-down of PDCD2L does. Because yeast Tsr4 does not bind nascent 40S and is essential for 40S production (32), PDCD2 may serve a role more similar to yeast Tsr4 than PDCD2L does. Both PDCD2L and PDCD2 also form extraribosomal heterotrimeric complexes with Rps2 and the methyltransferase PRMT3 (29). The formation of a homologous complex in yeast is unlikely because yeast Tsr4 and Hmt1 (Rmt1), the yeast homolog of PRMT3, appear to be spatially separated as Hmt1 is found in the nucleus (47), however we have not tested their interaction. PRMT3 and Hmt1 methylate two arginine residues in Rps2 (48–51), but their modification is substoichiometric (49, 52), and mutation of these residues to lysine or alanine in yeast Rps2 had no obvious effect on growth (data not shown) suggesting that their methylation is unimportant for basic cellular function and viability. In a more recent study, the Bachand group showed that PRMT3 does not co-translationally associate with Rps2 (53), but whether or not PDCD2L or PDCD2 co-translationally associate with Rps2 was not explored. Interestingly, the Bachand study showed that the human zinc-finger protein ZNF277 co-translationally binds to Rps2, however knock-down of ZNF277 does not impair 40S production, suggesting ZNF277 has a role outside of ribosome biogenesis (53). Higher eukaryotic organisms appear to have evolved a more elaborate mechanism for handling Rps2 compared to that of *S. cerevisiae*.

## Materials and Methods

### Strains, growth media, and genetic methods

All *S. cerevisiae* strains and sources are listed in Table 1. AJY2873 was generated by integration of a fragment of pAJ1915 to delete nucleotides 190 to 266 of URA3 in the *tsr4-ts* strain obtained from (54). AJY3398 was generated by genomic integration of *KanMX6-P*_*GAL1*_ into the genomic locus of *RPS2* in BY4741. AJY4297 and AJY4300 were generated by integration *KanMX6-P*_*GAL1*_*-3HA* into the genomic locus of *TSR4* (55) and *KanMX6-P*_*GAL1*_ (55) into the genomic locus of *RPS2* in CH1305 (56), respectively, and the *XRN1* locus in the resultant strains was disrupted with *URA3*. AJY4298 was generated by integration *KanMX6-P*_*GAL1*_*-3HA* (55) into the genomic locus of *TSR4* in BY4741. AJY4353 was generated by integration of *TAP-HIS3MX6* into the genomic locus of *TSR4* in BY4741. AJY4359 was generated by disruption of the genomic locus of *TSR4* with integration of *KanMX6* (55) in BY4743 (Open Biosystems). The resultant strain was transformed with pAJ4183 and was subsequently sporulated and dissected. AJY4352 was generated by integration of TEV-13myc::*Nat*^*R*^ into the genomic locus of *TSR4* in BY4741. AJY4363 was generated by genomic integration of *TSR4-TEV-13myc::Nat*^*R*^ amplified from AJY4352 into AJY1134. All yeast were cultured at 30°C in either YPD (2% peptone, 1% yeast extract, 2% dextrose), YPgal (2% peptone, 1% yeast extract, 1% galactose), or synthetic dropout (SD) medium containing 2% dextrose unless otherwise noted. When appropriate, media were supplemented with 150-250 µg/ml G418 or 100 µg/ml Nourseothricin. All plasmids and sources are listed in Table 2.

**Table 1:** Yeast strains used in this study.

| Strain | Genotype | Source |
| --- | --- | --- |
| AJY1134 | *MATa pep4-3 prb1-1122 ura3-52 leu2-3,112 reg1-501 gal1* | (33) |
| AJY1548 | *MATalpha CRM1(T539C)HA his3Δ1 leu2Δ0 ura3Δ0 met15Δ0* | (41) |
| AJY1951 | *MATa his3Δ1 leu2Δ0 met15Δ0 ura3Δ0 SQT1-TAP::HIS3MX6* | (65) |
| AJY2873 | *MATa ura3Δ0 leu2Δ0 his3Δ1 can1Δ::LEU2-MFA1pr::His3 tsr4-ts::ura3Δ* | This study. |
| AJY3398 | *MATa his3Δ1 leu2Δ0 met15Δ0 ura3Δ0 KanMX6::P<sub>GAL1</sub>-RPS2* | This study. |
| AJY4251 | *MATa TSR4-GFP::HIS3MX6 leu2Δ0 ura3Δ0* | (66) |
| AJY4297 | *MATa ade2 ade3 lys2-801 ura3-52 KanMX6::P<sub>GAL1</sub>-3HA-TSR4 xrn1Δ::URA3* | This study. |
| AJY4298 | *MATa his3Δ1 leu2Δ0 met15Δ0 ura3Δ0 KanMX6::P<sub>GAL1</sub>-3HA-TSR4* | This study. |
| AJY4300 | *MATa ade2 ade3 lys2-801 ura3-52 KanMX6::P<sub>GAL1</sub>-RPS2 xrn1Δ::URA3* | This study. |
| AJY4352 | *MATa his3Δ1 leu2Δ0 met15Δ0 ura3Δ0 TSR4-TEV-13myc::Nat<sup>R</sup>* | This study. |
| AJY4353 | *MATa his3Δ1 leu2Δ0 met15Δ0 ura3Δ0 TSR4-TAP::HIS3MX6* | This study. |
| AJY4359 | *MATa ura3Δ0 leu2Δ0 his3Δ1 tsr4Δ::KanMX/pAJ4183 (TSR4 URA3 CEN ARS)* | This study. |
| AJY4363 | *MATa pep4-3 prb1-1122 ura3-52 leu2-3,112 reg1-501 gal1 TSR4-TEV-13myc::Nat<sup>R</sup>* | This study. |
| BY4741 | *MATa his3Δ1 leu2Δ0 met15Δ0 ura3Δ0* | Open Biosystems |
| RKY1977 | *MATa ade2 ade3 leu2 lys2-801 ura3-52 xrn1Δ* | (67) |

**Table 2:** Plasmids used in this study.

| Plasmid | Description | Source |
| --- | --- | --- |
| pAJ1399 | *RPS2-GFP HIS3 CEN ARS* | (68) |
| pAJ3576 | *RPS2<sub>1-73</sub>-GFP HIS3 CEN ARS* | This study |
| pAJ3577 | *RPS2<sub>74-254</sub>-GFP HIS3 CEN ARS* | This study |
| pAJ3578 | *RPS2<sub>1-223</sub>-GFP HIS3 CEN ARS* | This study |
| pAJ4081 | *P<sub>GAL1</sub>-RPS2<sub>1-73</sub>-GFP URA3 CEN ARS* | This study |
| pAJ4082 | *P<sub>GAL1</sub>-RPS2<sub>1-223</sub>-GFP URA3 CEN ARS* | This study |
| pAJ4183 | *TSR4 URA3 CEN ARS* | This study |
| pAJ4184 | *TSR4 URA3 2μ ARS* | This study |
| pAJ4186 | *TSR4-GFP URA3 CEN ARS* | This study |
| pAJ4207 | *RPS2 HIS3 CEN ARS* | This study |
| pAJ4219 | *P<sub>GAL1</sub>-RPS2<sub>1-73</sub>-GFP<sub>ΔBsrGI</sub> URA3 CEN ARS* | This study |
| pAJ4220 | *RPS2<sub>1-235</sub>-GFP HIS3 CEN ARS* | This study |
| pAJ4231 | *RPS2<sub>Δ74-223</sub>-GFP HIS3 CEN ARS* | This study |
| pAJ4232 | *TSR4 HIS3 CEN ARS* | This study |
| pAJ4233 | *RPS2<sub>74-223</sub>-GFP HIS3 CEN ARS* | This study |
| pAJ4234 | *RPS2<sub>224-254</sub>-GFP HIS3 CEN ARS* | This study |
| pAJ4241 | *P<sub>GAL1</sub>-RPS2<sub>Δ74-223</sub>-GFP URA3 CEN ARS* | This study |
| pAJ4242 | *P<sub>GAL1</sub>- RPS2<sub>74-223</sub>-GFP URA3 CEN ARS* | This study |
| pAJ4243 | *P<sub>GAL1</sub>- RPS2<sub>224-254</sub>-GFP URA3 CEN ARS* | This study |
| pAJ4244 | *RPS2 URA3 CEN ARS* | This study |
| pAJ4261 | *P<sub>GAL1</sub>-RPS2-GFP URA3 CEN ARS* | This study |
| pAJ4267 | *TSR4<sub>ΔNLS</sub>-GFP URA3 CEN ARS* | This study |
| pAJ4270 | *P<sub>GAL1</sub> -GFP URA3 CEN ARS* | This study |
| pAJ4271 | *P<sub>GAL1</sub>-RPS2<sub>74-254</sub>-GFP URA3 CEN ARS* | This study |
| pAJ4451 | *P<sub>GAL1</sub>-RPS2<sub>1-73</sub>_F14S-GFP URA3 CEN ARS* | This study |
| pAJ4452 | *P<sub>GAL1</sub>-RPS2<sub>1-73</sub>_G15D-GFP URA3 CEN ARS* | This study |
| pAJ4453 | *P<sub>GAL1</sub>-RPS2<sub>1-73</sub>_N18D-GFP URA3 CEN ARS* | This study |
| pAJ4454 | *P<sub>GAL1</sub>-RPS2<sub>1-73</sub>_W35R-GFP URA3 CEN ARS* | This study |
| pAJ4455 | *P<sub>GAL1</sub>-RPS2<sub>1-73</sub>_P37L-GFP URA3 CEN ARS* | This study |
| pAJ4456 | *P<sub>GAL1</sub>-RPS2<sub>1-73</sub>_V38D-GFP URA3 CEN ARS* | This study |
| pAJ4457 | *P<sub>GAL1</sub>-RPS2<sub>1-73</sub>_R43G-GFP URA3 CEN ARS* | This study |
| pAJ4458 | *P<sub>GAL1</sub>-RPS2<sub>1-73</sub>_H59R-GFP URA3 CEN ARS* | This study |
| pAJ4459 | *P<sub>GAL1</sub>-RPS2<sub>1-73_L61S</sub>-GFP URA3 CEN ARS* | This study |
| pAJ4460 | *P<sub>GAL1</sub>-RPS2<sub>1-73_E65K</sub>-GFP URA3 CEN ARS* | This study |
| pRS413 | *HIS3 CEN ARS* | (69) |
| pRS416 | *URA3 CEN ARS* | (69) |
| pRS426 | *URA3 2μ ARS* | (69) |

### Random PCR mutagenesis of Rps2_1-73_-GFP

Random mutations in *P*_*GAL1*_*-RPS2*_*1-73*_*-GFP*, were generated by error-prone PCR using Taq polymerase and pAJ4081 as template with oligos that hybridize to sequence upstream of the *GAL1* promoter and to the *GFP* coding sequence. The vector pAJ4219 was linearized with the restriction enzymes *Bsr*GI and *Bam*HI and co-transformed with the mutant amplicon library into BY4741 and plated onto SD-ura media containing glucose to allow gap rescue of the linearized vector via the promoter and *GFP* sequences. Approximately 3000 colonies were screened on SD-ura containing galactose for their ability to alleviate the toxicity caused by the expression of the Rps2_1-73_-GFP fragment. Vectors expressing mutants with reduced toxicity and that showed GFP expression were rescued from yeast, transformed back into BY4741 to verify their alleviation of the toxicity and sequenced.

### Immunoprecipitation

For purifying the panel of Rps2-GFP fragments, strain AJY1134 containing the appropriate galactose-inducible vectors was cultured in glucose-containing media until exponential phase. Expression was then induced for 90 min by the addition of galactose to 1% final concentration. Subsequent steps were performed on ice or at 4°C unless otherwise noted. Cells were washed once in Buffer A (20 mM HEPES-KOH pH8.0, 110 mM KOAc, 40 mM NaCl, 1mM PMSF and benzamidine, and 1 μM leupeptin and pepstatin), resuspended in Buffer A and extracts were prepared by vortexing with glass beads. Extracts were clarified by centrifugation at 18,213 x *g* for 15 min, normalized to A_260_ units and TritonX-100 was added to a final concentration of 0.1% (v/v). The extracts were subjected to ultracentrifugation at 70 krpm for 15 min on a TLA100.3 rotor (Beckman-Coulter) to remove ribosomes. Supernatants were incubated with GFP-trap (Chromotek) for 90 minutes. The beads were washed three times with Buffer A-w (Buffer A supplemented with 0.1% (v/v) TritonX-100), resuspended in 1X Laemmli buffer and heated at 99°C for 3 minutes. Samples were separated on 6-18% SDS-PAGE gradient gels, bands of interest were excised. Protein-containing gel slices were prepared for in-gel digestion and peptides recovered for identification by mass spectrometry as described (57). The resultant peptides were run for 30 minutes on a Dionex LC and Orbitrap Fusion 1 (Thermo Scientific) for LC-MS/MS. Data were processed in Scaffold v4.8.3 (Proteome Software, Inc.), and a protein threshold of 99% minimum and 2 peptides minimum, and peptide threshold of 0.5% FDR was applied.

To assess the interactions between Tsr4-TEV-13myc and the Rps2-GFP fragments or between Tsr4-TEV-13myc and the Rps2_1-73_-GFP mutants, we transformed the appropriate galactose-inducible vectors into AJY4363. Cells were cultured and extracts prepared as described above without ribosome removal and immunoprecipitations were done with anti-*c*-Myc magnetic beads (Pierce 88842). IPs and Inputs were separated on 6-18% SDS-PAGE gradient gels and subjected to Western Blotting (see below).

### Sucrose density gradient analysis

AJY3398 was transformed with pAJ1399, pAJ3576, pAJ3578, and pRS413. Cells were cultured in glucose-containing media lacking histidine for 5 hours at 30°C to deplete endogenous Rps2 then treated with 100 µg/mL CHX for 10 minutes at 30°C. Extracts and sucrose density gradient fractionation was done as previously described (57) except the lysis buffer contained 100 µg/mL of CHX and 7 mM of beta-mercaptoethanol.

### Western blotting

Primary antibodies used in this study were anti-*c-*Myc monoclonal 9e10 (Biolgend), anti-GFP (M. Rout), anti-Rpl8 (K-Y. Lo), anti-Rps24 (our lab), and anti-Xrn1 (our lab). Secondary antibodies used were goat anti-mouse IRDye 800CW (LI-COR Biosciences) and goat anti-rabbit IRDye 680RD (LI-COR Biosciences). Blots were imaged with an Odyssey CLx Infrared Imaging System (LI-COR Biosciences) using Image Studio (LI-COR Biosciences).

### Assay for co-translational association by immunoprecipitation and RT-qPCR

For strains expressing the TAP-tagged bait proteins, cultures of AJY1951 and AJY4353 were grown to exponential phase then treated with 100 µg/ml of CHX for 10 minutes prior to harvesting. After harvesting, cells were washed with Buffer B (50 mM Tris pH 7.5, 100 mM NaCl, 1.5 mM MgCl2, 100 µg/mL CHX, 1mM PMSF and benzamidine, and 1 μM leupeptin and pepstatin), resuspended in the same buffer and extracts prepared by vortexing with glass beads. Extracts were clarified by centrifugation at 18,213 x *g* for 10 min, normalized by A_260_ units and TritonX-100 was added to a final concentration of 0.1% (v/v). 3% of each normalized extract were taken as Input. The normalized extracts were incubated with Dynabeads (Invitrogen) coupled to rabbit IgG (Sigma), prepared as described (58), for 90 minutes. The beads were washed three times with Buffer B-w (Buffer B supplemented with 0.1% TritonX-100). RNAs from the IPs and Inputs were isolated using acid-phenol-chloroform as described (59) and were resuspended in nuclease-free water. For GFP-tagged Tsr4, AJY4251 was cultured and affinity purification was done similarly except extracts were generated in Buffer A (see “Immunoprecipitation” section above) and bead washes used Buffer A-w both supplemented with 100 µg/mL CHX, and protein-G Dynabeads (Invitrogen) precoated with rabbit anti-GFP (M. Rout) were used for immunoprecipitation.

Fifty nanograms of RNA from each IP and Input was used as template for cDNA synthesis using qScript cDNA SuperMix kit (Quantabio 95048-025). For qPCR, 10% (for the TAP-tagged samples) and 13% (for the GFP-tagged sample) of each reverse transcription (RT) reaction was used as template using PerfeCTa SYBR Green FastMix Low ROX kit (Quantabio 95074-250). qPCR for each oligonucleotide pair was done in technical triplicate. The following program on an Applied Biosystems ViiA 7 Real-Time PCR instrument was used: an initial hold step at 50°C for 2 minutes followed by an initial denaturation and polymerase activation step at 95°C for 10 minutes, followed by 40 cycles of denaturation at 95°C for 15 s and elongation and data collection at 60°C for 1 minute. A ramp rate of 1°C/s between thermocycling was used. The following oligonucleotide pairs were used: *RPS2*-Forward 5’-GAAGATGTCTACACCCAATCTAACG-3’ and *RPS2*-Reverse 5’-GAGTCAAGAAACCGTATGTGTTACC-3’ (amplicon size of 100 bp), and *RPL10*-Forward 5’-AGATACCAAAAGAACAAGCCTTACC-3’ and *RPL10*-Reverse 5’-GTAGATTCTGATCTTGGAGTCTGGA-3’ (amplicon size of 75 bp). Oligonucleotide pairs were designed using the Primer3web software version 4.1.0 (http://primer3.ut.ee).

The threshold cycle (C_t_) for each reaction was incorporated into the standard curve to derive the number of mRNA molecules present using the ViiA 7 software (Applied Biosystems). The standard curve was generated using 13, 1.3, and 0.13 ng of genomic DNA from BY4741 corresponding to about 1,000,000, 100,000, and 10,000 genomic loci for each target gene. The number of RNA molecules isolated in the whole IP (IP_RNA_) and the number of molecules present in the entire Input (Input_RNA_) were extrapolated. The ratio of IP_RNA_ to Input_RNA_ was calculated for each reaction. The average ratio and standard deviation for each triplicate set was determined. The ratios were then normalized to the average ratio of the target mRNA for each bait to display enrichment relative to the target mRNA and plotted in Graphpad Prism version 7.0c.169 for Mac iOS (https://www.graphpad.com).

### Rps2-GFP expression in *tsr4-ts* strain

Strains AJY2873 and BY4741 containing pAJ4261 were grown to mid log phase in 100 mL of SD-ura medium containing raffinose and then shifted to 37°C for 2 hours. Rps2-GFP expression was induced for 90 min with 1% galactose prior to harvesting the cells. All subsequent steps were carried out on ice or at 4°C. Cells were washed and resuspended in Buffer A, extracts were made by glass bead lysis, clarified for 10 minutes at 18,213 x *g* and normalized to 1 A_260_ unit in a final volume of 100 µL. Half of each normalized extract was overlaid onto a 50 µL 15% sucrose cushion in the same buffer and subjected to ultracentrifugation at 70 krpm for 15 minutes on a TLA100 rotor. Fractions were taken by manual pipetting, and Laemmli sample buffer was added to 1X concentration. The samples were heated at 99°C for 3 minutes. Equal relative amounts of each sample were separated on a 6-18% SDS-PAGE gradient gel and subjected to analysis by Western blotting.

### Fluorescent *in situ* hybridization (FISH)

RKY1977, AJY4297 and AJY4300 cells were grown to saturation in galactose-containing media, diluted 5-fold in fresh glucose-containing media and continued to grow for 2 hours. Formaldehyde was added to 4.5% final concentration and cultures gently agitated at 30°C for 30 mi. Cells were washed twice with KSorb buffer (1.2 M Sorbitol, 0.1 M Potassium phosphate buffer 7.0) and then incubated in KSorb buffer with 50 µg/ml Zymolyase T20 for 15 minutes at 37°C in presence of 20 mM Vanadyl Ribonucloside complex (VRC), 28 mM β-mercaptoethanol and 1 mM PMSF. Cells were gently pelleted, washed three times with ice cold KSorb buffer and resuspended in Ksorb buffer before applying to Teflon-coated Immunofluorescence slides (Polysciences Inc., No. 18357) pre-coated with Poly-lysine. Slides were incubated in a moist chamber at room temperature for 10 minutes, excess cells were gently aspirated, and the slides were stored in 70% ethanol at −20°C. Cells were rehydrated by washing twice with 2X SSC (300 mM NaCl, 30 mM Sodium Citrate pH 7.0) and then incubated in Prehybridization solution (10% Dextran sulfate, 50% deionized formamide, 1X Denhardt’s, 2 mM VRC and 4X SSC, 0.2% purified BSA, 25 µg yeast tRNA and 500 µg/ml ssDNA) for 1h at 72°C in a moist chamber. Solution was replaced with Prehybridization solution containing 1 µM Cy3-labelled oligonucleotide D-A2 probe (5’-ATGCTCTTGCCAAAACAAAAAAATCCATTTTCAAAATTATTAAATTTCTT-3’). Slides were incubated in a moist chamber at 72°C for 1h followed by overnight incubation at 37°C. Wells were then washed with 2X SSC at 37°C and then 1X SSC at room temperature containing 0.1% NP-40 for 30 minutes each. Cells were incubated for 2 minutes with 4’,6’-diamidino-2-phenylindole (DAPI) at 1 µg/ml in PBS, washed twice with PBS and mounted in Aqua-Poly/Mount (Polysciences Inc.). Fluorescence was visualized on a Nikon E800 microscope fitted with a Plan Apo 100×/1.4 objective and a Photometrics CoolSNAP ES camera controlled by NIS-Elements AR2.10 software and photos were processed with Affinity Designer.

### Fluorescent microscopy of GFP-tagged proteins

Cells were cultured as described in the figure legends. Fluorescent signal was captured using a 500 ms exposure time unless otherwise noted on a Nikon E800 microscope fitted with a Plan Apo 100×/1.4 objective and a Photometrics CoolSNAP ES camera controlled by NIS-Elements AR2.10 software and photos were processed with Affinity Designer.

## Acknowledgements

This work was supported by grants NIH GM108823 and GM127127 to AWJ and by a fellowship from the University of Texas at Austin Graduate School to JJB. The Proteomics Facility in the Center for Biomedical Research Support at the University of Texas at Austin is supported in part by the CPRIT grant RP110782. We thank M. Rout and K-Y. Lo for antibodies. We also wish to thank K. Robbins and J. Recchia-Rife for help with experiments and L. Leblanc for advice regarding RT-qPCR. We thank P. Sutjita for helpful comments regarding the manuscript.

## Author Contributions

JJB and AWJ conceptualized and performed experiments, analyzed and interpreted data, and composed and edited the manuscript and figures. SM performed the FISH analysis and interpreted the data.

---

**Figure S1.**
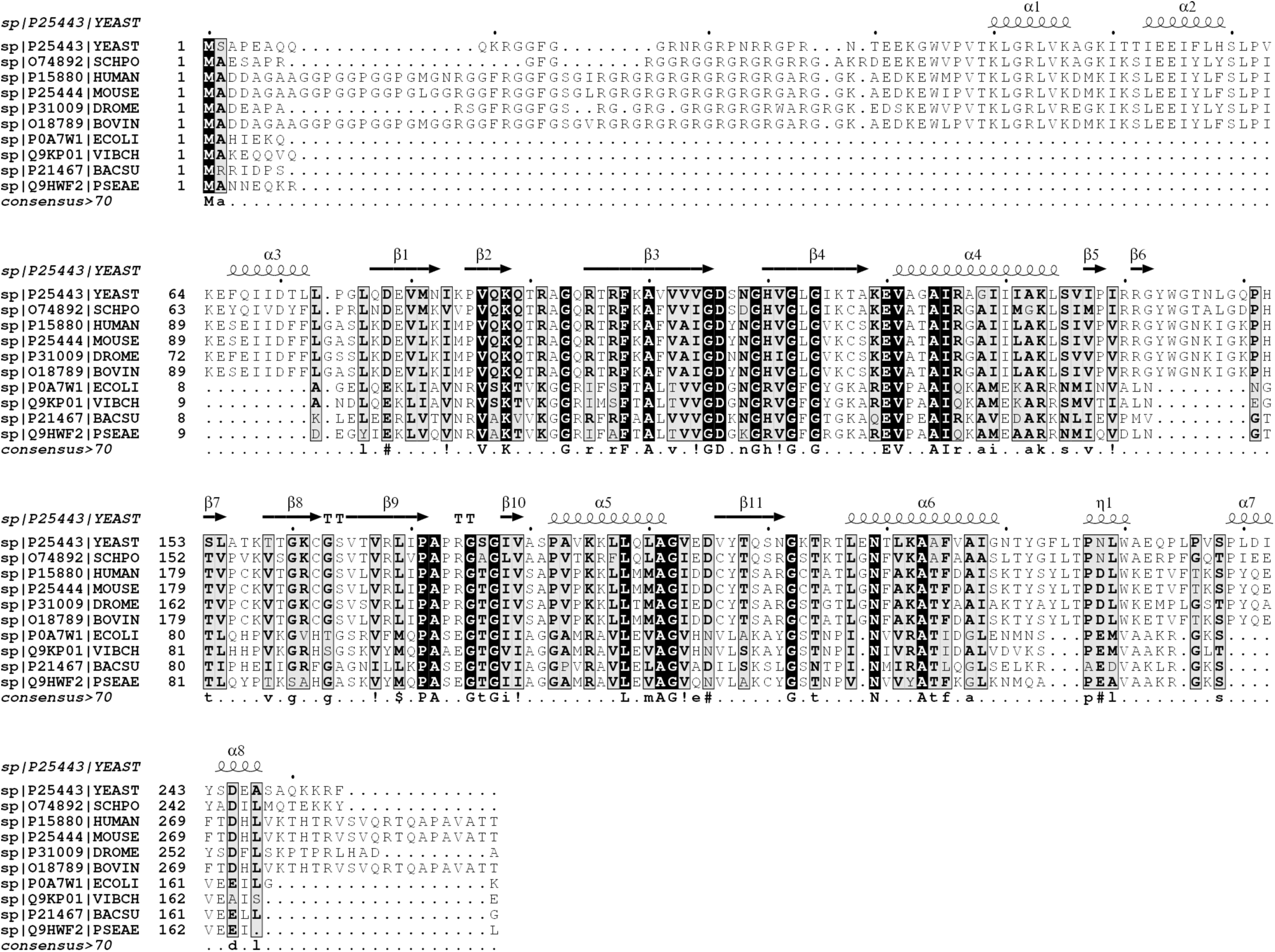
Multiple sequence alignment of Rps2 from selected eukaryotic and prokaryotic species. Multiple sequence alignment of Rps2 orthologs from various species using the T-Coffee web server (references (60, 61) in the main text). The secondary structure of yeast Rps2 ((27) in the main text) was fitted above the alignment using ESPript 3.0 web server ((62) in the main text). Residues are shaded as a percentage of similarity considering physico-chemical properties. Helices are shown as squiggles, beta sheets are shown as arrows, and turns are indicated by “TT”. Species are annotated as: YEAST (*S. cerevisiae*), SCHPO (*S. pombe*), HUMAN (*H. sapiens*), MOUSE (*M. musculus*), DROME (*D. melanogaster*), BOVIN (*B. taurus*), ECOLI, (*E. coli*), VIBCH (*V. cholera*), BACSU (*B. subtilis*), and PSAEE (*P. aeruginosa*).

**Figure S2.**
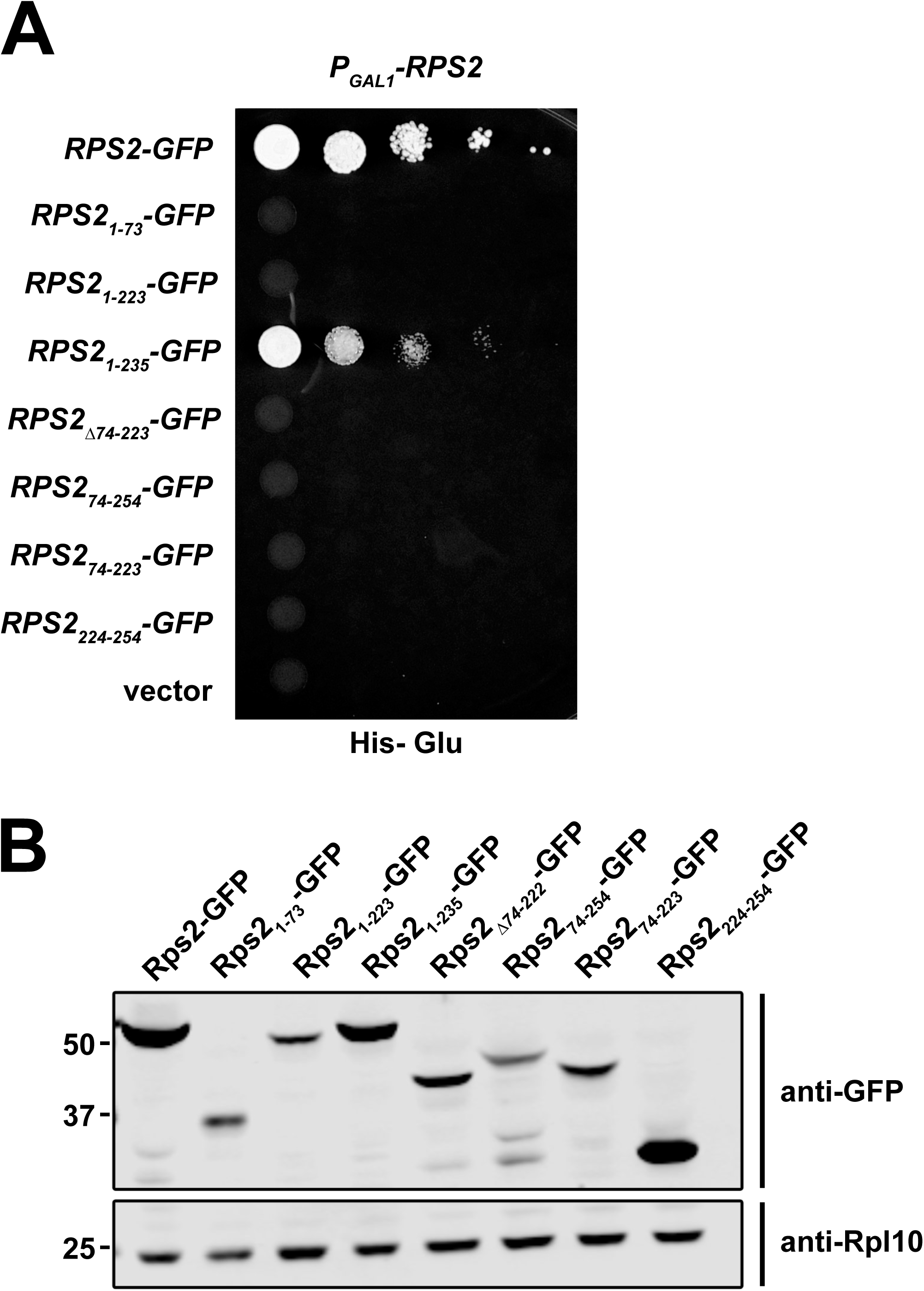
Complementation and expression of the Rps2-GFP fragments. A) Growth assay of *P*_*GAL1*_*-RPS2* cells (AJY3398) with vectors expressing the indicated Rps2-GFP fragments under control of the native *RPS2* promoter as shown by 10-fold serial dilutions plated on glucose-containing media lacking histidine grown for 2 days. B) The expression of the Rps2-GFP fragments under the transcriptional control of their native promoter from centromeric vectors in BY4741 grown to late exponential phase in glucose-containing media lacking histidine. Rpl10 was used as a loading control. Proteins were extracted as in ((63) in the main text), separated by SDS-PAGE and analyzed by Western blotting.

**Figure S3.**
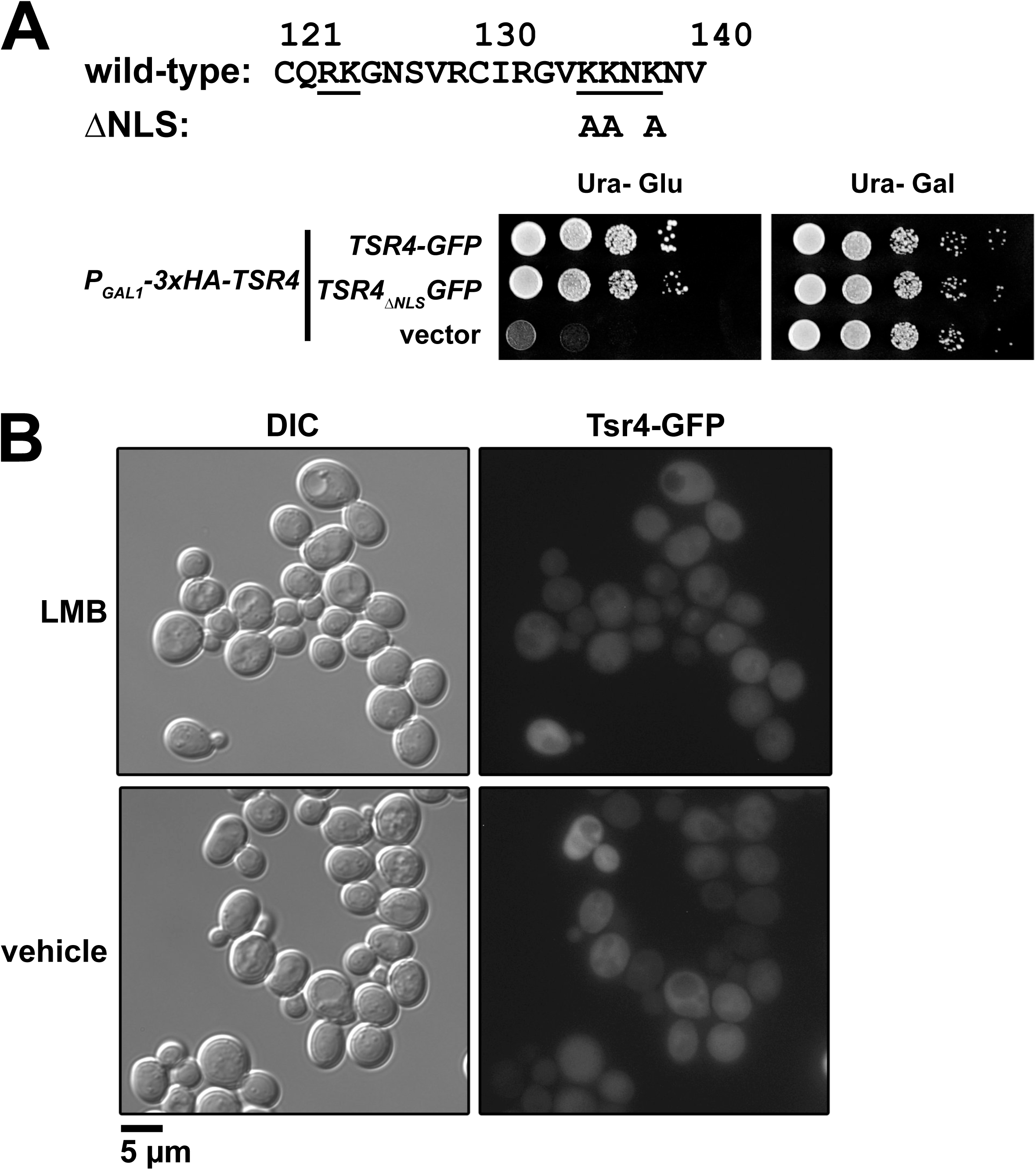
Tsr4 does not shuttle between the nucleus and cytoplasm. A) The growth of *P*_*GAL1*_*-3xHA-TSR4* (AJY4298) cells harboring empty vector (pRS416) or vectors encoding *TSR4-GFP* (pAJ4186) or *TSR4*_*ΔNLS*_*-GFP* (pAJ4267) as shown by 10-fold serial dilutions plated on glucose-containing media lacking uracil after 2 days. Primary sequence for residues 121-140 of Tsr4 is shown above. The putative bi-partite NLS predicted by PSORT II ((64) in the main text) is underlined; alanine substitutions of Tsr4_ΔNLS_ are indicated below the sequence. B) Subcellular localization of Tsr4-GFP (pAJ4186) after 30 min inhibition of Crm1_T539C_ by LMB in strain AJY1548. Treatment with 1% ethanol (EtOH) was used in parallel as a vehicle control.

**Figure S4.**
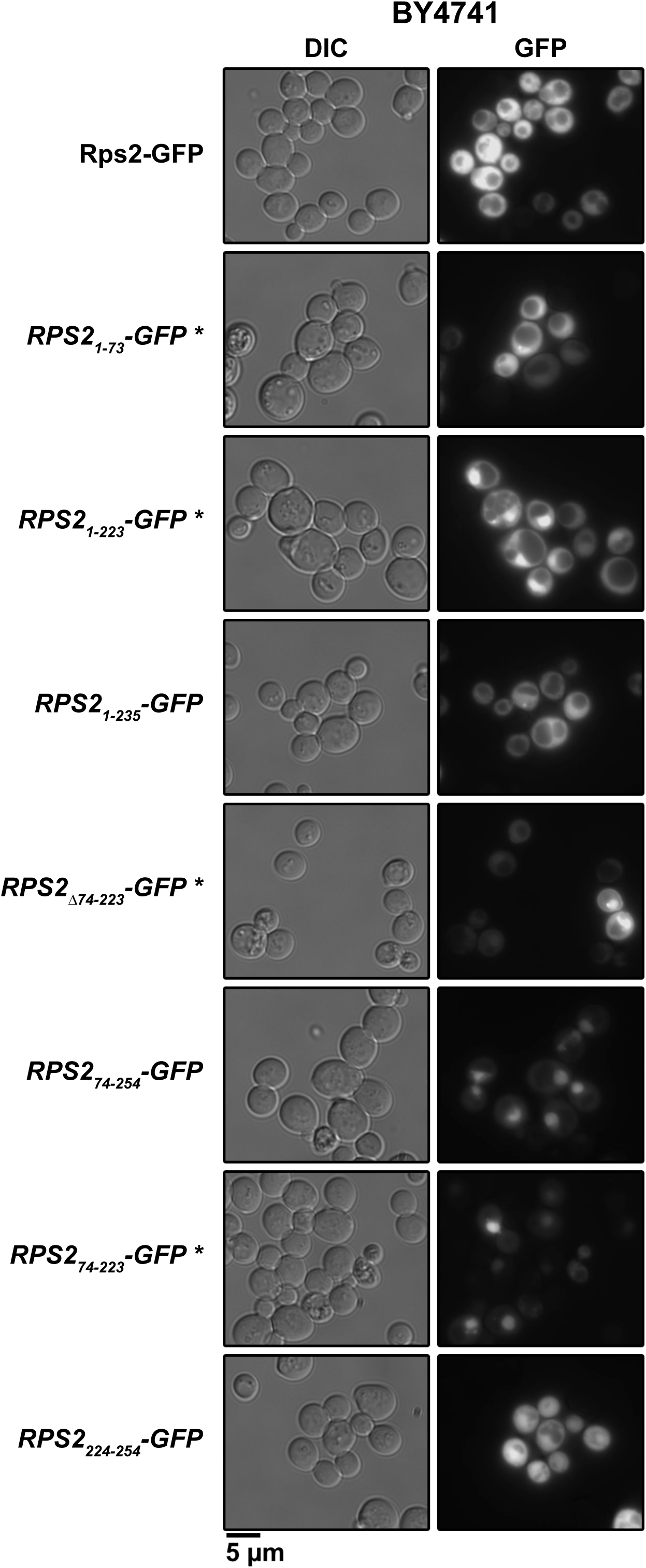
Subcellular localization of Rps2-GFP fragments. Steady-state distribution of the Rps2-GFP fragments expressed from their native promoter from centromeric vectors in BY4741 cells as analyzed by fluorescent microscopy. Cells were from the same cultures as the protein extracts shown in Fig. S2B. A 500ms exposure time was used to capture GFP signal except for the samples marked with an asterisks (*) which were captured using a 2s exposure time.

## References

1. Woolford JL, Baserga SJ. 2013. Ribosome biogenesis in the yeast *Saccharomyces cerevisiae*. Genetics 195:643–81.

2. Kressler D, Hurt E, Baßler J. 2017. A Puzzle of Life: Crafting Ribosomal Subunits. Trends Biochem Sci 42:640–654.

3. Peña C, Hurt E, Panse VG. 2017. Eukaryotic ribosome assembly, transport and quality control. Nat Struct Mol Biol 24:689–699.

4. Baßler J, Hurt E. 2018. Eurokaryotic Ribosome Assembly. Annu Rev Biochem 88:8.1–8.26.

5. Jakob S, Ohmayer U, Neueder A, Hierlmeier T, Perez-Fernandez J, Hochmuth E, Deutzmann R, Griesenbeck J, Tschochner H, Milkereit P. 2012. Interrelationships between yeast ribosomal protein assembly events and transient ribosome biogenesis factors interactions in early pre-ribosomes. PLoS One 7:e32552.

6. Ferreira-Cerca S, Pöll G, Gleizes P-E, Tschochner H, Milkereit P. 2005. Roles of eukaryotic ribosomal proteins in maturation and transport of pre-18S rRNA and ribosome function. Mol Cell 20:263–75.

7. Pöll G, Braun T, Jakovljevic J, Neueder A, Jakob S, Woolford JL, Tschochner H, Milkereit P. 2009. rRNA maturation in yeast cells depleted of large ribosomal subunit proteins. PLoS One 4:e8249.

8. Jäkel S, Mingot J-M, Schwarzmaier P, Hartmann E, Görlich D. 2002. Importins fulfil a dual function as nuclear import receptors and cytoplasmic chaperones for exposed basic domains. EMBO J 21:377–86.

9. Pillet B, Mitterer V, Kressler D, Pertschy B. 2017. Hold on to your friends: Dedicated chaperones of ribosomal proteins: Dedicated chaperones mediate the safe transfer of ribosomal proteins to their site of pre-ribosome incorporation. Bioessays 39:1–12.

10. Ting Y-H, Lu T-J, Johnson AW, Shie J-T, Chen B-R, Kumar S S, Lo K-Y. 2017. Bcp1 Is the Nuclear Chaperone of Rpl23 in Saccharomyces cerevisiae. J Biol Chem 292:585–596.

11. Eisinger DP, Dick FA, Denke E, Trumpower BL. 1997. SQT1, which encodes an essential WD domain protein of Saccharomyces cerevisiae, suppresses dominant-negative mutations of the ribosomal protein gene QSR1. Mol Cell Biol 17:5146–55.

12. Schaper S, Fromont-Racine M, Linder P, de la Cruz J, Namane A, Yaniv M. 2001. A yeast homolog of chromatin assembly factor 1 is involved in early ribosome assembly. Curr Biol 11:1885–90.

13. Iouk TL, Aitchison JD, Maguire S, Wozniak RW. 2001. Rrb1p, a yeast nuclear WD-repeat protein involved in the regulation of ribosome biosynthesis. Mol Cell Biol 21:1260–71.

14. Pausch P, Singh U, Ahmed YL, Pillet B, Murat G, Altegoer F, Stier G, Thoms M, Hurt E, Sinning I, Bange G, Kressler D. 2015. Co-translational capturing of nascent ribosomal proteins by their dedicated chaperones. Nat Commun 6:7494.

15. Pillet B, García-Gómez JJ, Pausch P, Falquet L, Bange G, de la Cruz J, Kressler D. 2015. The Dedicated Chaperone Acl4 Escorts Ribosomal Protein Rpl4 to Its Nuclear Pre-60S Assembly Site. PLoS Genet 11:e1005565.

16. Stelter P, Huber FM, Kunze R, Flemming D, Hoelz A, Hurt E. 2015. Coordinated Ribosomal L4 Protein Assembly into the Pre-Ribosome Is Regulated by Its Eukaryote-Specific Extension. Mol Cell 58:854–62.

17. Koch B, Mitterer V, Niederhauser J, Stanborough T, Murat G, Rechberger G, Bergler H, Kressler D, Pertschy B. 2012. Yar1 protects the ribosomal protein Rps3 from aggregation. J Biol Chem 287:21806–15.

18. Kressler D, Bange G, Ogawa Y, Stjepanovic G, Bradatsch B, Pratte D, Amlacher S, Strauß D, Yoneda Y, Katahira J, Sinning I, Hurt E. 2012. Synchronizing nuclear import of ribosomal proteins with ribosome assembly. Science 338:666–71.

19. Holzer S, Ban N, Klinge S. 2013. Crystal structure of the yeast ribosomal protein rpS3 in complex with its chaperone Yar1. J Mol Biol 425:4154–60.

20. Huber FM, Hoelz A. 2017. Molecular basis for protection of ribosomal protein L4 from cellular degradation. Nat Commun 8:14354.

21. West M, Hedges JB, Chen A, Johnson AW. 2005. Defining the order in which Nmd3p and Rpl10p load onto nascent 60S ribosomal subunits. Mol Cell Biol 25:3802–13.

22. Rout MP, Blobel G, Aitchison JD. 1997. A distinct nuclear import pathway used by ribosomal proteins. Cell 89:715–25.

23. Mitterer V, Murat G, Réty S, Blaud M, Delbos L, Stanborough T, Bergler H, Leulliot N, Kressler D, Pertschy B. 2016. Sequential domain assembly of ribosomal protein S3 drives 40S subunit maturation. Nat Commun 7:10336.

24. Mitterer V, Gantenbein N, Birner-Gruenberger R, Murat G, Bergler H, Kressler D, Pertschy B. 2016. Nuclear import of dimerized ribosomal protein Rps3 in complex with its chaperone Yar1. Sci Rep 6:36714.

25. Calviño FR, Kharde S, Ori A, Hendricks A, Wild K, Kressler D, Bange G, Hurt E, Beck M, Sinning I. 2015. Symportin 1 chaperones 5S RNP assembly during ribosome biogenesis by occupying an essential rRNA-binding site. Nat Commun 6:6510.

26. Bange G, Murat G, Sinning I, Hurt E, Kressler D. 2013. New twist to nuclear import: When two travel together. Commun Integr Biol 6:e24792.

27. Ben-Shem A, Garreau de Loubresse N, Melnikov S, Jenner L, Yusupova G, Yusupov M. 2011. The structure of the eukaryotic ribosome at 3.0 Å resolution. Science 334:1524–9.

28. Perreault A, Bellemer C, Bachand F. 2008. Nuclear export competence of pre-40S subunits in fission yeast requires the ribosomal protein Rps2. Nucleic Acids Res 36:6132–42.

29. Landry-Voyer A-M, Bilodeau S, Bergeron D, Dionne KL, Port SA, Rouleau C, Boisvert F-M, Kehlenbach RH, Bachand F. 2016. Human PDCD2L Is an Export Substrate of CRM1 That Associates with 40S Ribosomal Subunit Precursors. Mol Cell Biol 36:3019–3032.

30. Minakhina S, Naryshkina T, Changela N, Tan W, Steward R. 2016. Zfrp8/PDCD2 Interacts with RpS2 Connecting Ribosome Maturation and Gene-Specific Translation. PLoS One 11:e0147631.

31. Burroughs AM, Aravind L. 2014. Analysis of two domains with novel RNA-processing activities throws light on the complex evolution of ribosomal RNA biogenesis. Front Genet 5:424.

32. Li Z, Lee I, Moradi E, Hung N-J, Johnson AW, Marcotte EM. 2009. Rational extension of the ribosome biogenesis pathway using network-guided genetics. PLoS Biol 7:e1000213.

33. Hovland P, Flick J, Johnston M, Sclafani RA. 1989. Galactose as a gratuitous inducer of GAL gene expression in yeasts growing on glucose. Gene 83:57–64.

34. Ho B, Baryshnikova A, Brown GW. 2018. Unification of Protein Abundance Datasets Yields a Quantitative Saccharomyces cerevisiae Proteome. Cell Syst 6:192–205.e3.

35. Castello A, Fischer B, Eichelbaum K, Horos R, Beckmann BM, Strein C, Davey NE, Humphreys DT, Preiss T, Steinmetz LM, Krijgsveld J, Hentze MW. 2012. Insights into RNA biology from an atlas of mammalian mRNA-binding proteins. Cell 149:1393–406.

36. Fatica A, Oeffinger M, Dlakić M, Tollervey D. 2003. Nob1p is required for cleavage of the 3’ end of 18S rRNA. Mol Cell Biol 23:1798–807.

37. Fatica A, Tollervey D, Dlakić M. 2004. PIN domain of Nob1p is required for D-site cleavage in 20S pre-rRNA. RNA 10:1698–701.

38. Stevens A, Hsu CL, Isham KR, Larimer FW. 1991. Fragments of the internal transcribed spacer 1 of pre-rRNA accumulate in Saccharomyces cerevisiae lacking 5’ 3’ exoribonuclease 1. J Bacteriol 173:7024–8.

39. Moy TI, Silver PA. 1999. Nuclear export of the small ribosomal subunit requires the ran-GTPase cycle and certain nucleoporins. Genes Dev 13:2118–33.

40. Hutten S, Kehlenbach RH. 2007. CRM1-mediated nuclear export: to the pore and beyond. Trends Cell Biol 17:193–201.

41. Hedges J, West M, Johnson AW. 2005. Release of the export adapter, Nmd3p, from the 60S ribosomal subunit requires Rpl10p and the cytoplasmic GTPase Lsg1p. EMBO J 24:567–79.

42. Panasenko OO, Somasekharan SP, Villanyi Z, Zagatti M, Bezrukov F, Rashpa R, Cornut J, Iqbal J, Longis M, Carl SH, Peña C, Panse VG, Collart MA. 2019. Co-translational assembly of proteasome subunits in NOT1-containing assemblysomes. Nat Struct Mol Biol 26:1.

43. Collart MA, Kassem S, Villanyi Z. 2017. Mutations in the NOT Genes or in the Translation Machinery Similarly Display Increased Resistance to Histidine Starvation. Front Genet 8:1–7.

44. Sung M-K, Porras-Yakushi TR, Reitsma JM, Huber FM, Sweredoski MJ, Hoelz A, Hess S, Deshaies RJ. 2016. A conserved quality-control pathway that mediates degradation of unassembled ribosomal proteins. Elife 5:e19105.

45. Sung M-K, Reitsma JM, Sweredoski MJ, Hess S, Deshaies RJ. 2016. Ribosomal proteins produced in excess are degraded by the ubiquitin-proteasome system. Mol Biol Cell 27:2642–52.

46. Shiber A, Döring K, Friedrich U, Klann K, Merker D, Zedan M, Tippmann F, Kramer G, Bukau B. 2018. Cotranslational assembly of protein complexes in eukaryotes revealed by ribosome profiling. Nature 561:268–272.

47. Henry MF, Silver P a. 1996. A novel methyltransferase (Hmt1p) modifies poly(A)+-RNA-binding proteins. Mol Cell Biol 16:3668–78.

48. Yagoub D, Hart-Smith G, Moecking J, Erce MA, Wilkins MR. 2015. Yeast proteins Gar1p, Nop1p, Npl3p, Nsr1p, and Rps2p are natively methylated and are substrates of the arginine methyltransferase Hmt1p. Proteomics 15:3209–18.

49. Lipson RS, Webb KJ, Clarke SG. 2010. Rmt1 catalyzes zinc-finger independent arginine methylation of ribosomal protein Rps2 in Saccharomyces cerevisiae. Biochem Biophys Res Commun 391:1658–62.

50. Swiercz R, Cheng D, Kim D, Bedford MT. 2007. Ribosomal protein rpS2 is hypomethylated in PRMT3-deficient mice. J Biol Chem 282:16917–23.

51. Swiercz R, Person MD, Bedford MT. 2005. Ribosomal protein S2 is a substrate for mammalian PRMT3 (protein arginine methyltransferase 3). Biochem J 386:85–91.

52. Young BD, Weiss DI, Zurita-Lopez CI, Webb KJ, Clarke SG, McBride AE. 2012. Identification of methylated proteins in the yeast small ribosomal subunit: a role for SPOUT methyltransferases in protein arginine methylation. Biochemistry 51:5091–104.

53. Dionne KL, Bergeron D, Landry-Voyer A-M, Bachand F. 2018. The 40S ribosomal protein uS5 (RPS2) assembles into an extra-ribosomal complex with human ZNF277 that competes with the PRMT3-uS5 interaction. J Biol Chem 5:jbc.RA118.004928.

54. Ben-Aroya S, Coombes C, Kwok T, O’Donnell KA, Boeke JD, Hieter P. 2008. Toward a comprehensive temperature-sensitive mutant repository of the essential genes of Saccharomyces cerevisiae. Mol Cell 30:248–58.

55. Longtine MS, McKenzie A, Demarini DJ, Shah NG, Wach A, Brachat A, Philippsen P, Pringle JR. 1998. Additional modules for versatile and economical PCR-based gene deletion and modification in Saccharomyces cerevisiae. Yeast 14:953–61.

56. Kranz JE, Holm C. 1990. Cloning by function: an alternative approach for identifying yeast homologs of genes from other organisms. Proc Natl Acad Sci U S A 87:6629–33.

57. Black JJ, Wang Z, Goering LM, Johnson AW. 2018. Utp14 interaction with the Small Subunit Processome. RNA rna.066373.118.

58. Oeffinger M, Wei KE, Rogers R, DeGrasse JA, Chait BT, Aitchison JD, Rout MP. 2007. Comprehensive analysis of diverse ribonucleoprotein complexes. Nat Methods 4:951–6.

59. Zhu J, Liu X, Anjos M, Correll CC, Johnson AW. 2016. Utp14 Recruits and Activates the RNA Helicase Dhr1 To Undock U3 snoRNA from the Preribosome. Mol Cell Biol 36:965–78.

60. Notredame C, Higgins DG, Heringa J. 2000. T-Coffee: A novel method for fast and accurate multiple sequence alignment. J Mol Biol 302:205–17.

61. Di Tommaso P, Moretti S, Xenarios I, Orobitg M, Montanyola A, Chang J-M, Taly J-F, Notredame C. 2011. T-Coffee: a web server for the multiple sequence alignment of protein and RNA sequences using structural information and homology extension. Nucleic Acids Res 39:W13–7.

62. Robert X, Gouet P. 2014. Deciphering key features in protein structures with the new ENDscript server. Nucleic Acids Res 42:W320–4.

63. Kushnirov V V. 2000. Rapid and reliable protein extraction from yeast. Yeast 16:857–60.

64. Nakai K, Horton P. 1999. PSORT: a program for detecting sorting signals in proteins and predicting their subcellular localization. Trends Biochem Sci 24:34–6.

65. Ghaemmaghami S, Huh W-K, Bower K, Howson RW, Belle a, Dephoure N, O’Shea EK, Weissman JS. 2003. Global analysis of protein expression in yeast. Nature 425:737–741.

66. Huh W, Falvo J V, Gerke LC, Carroll AS, Howson RW, Weissman JS, O’Shea EK. 2003. Global analysis of protein localization in budding yeast. Nature 425:686–91.

67. Johnson AW, Kolodner RD. 1995. Synthetic lethality of sep1 (xrn1) ski2 and sep1 (xrn1) ski3 mutants of Saccharomyces cerevisiae is independent of killer virus and suggests a general role for these genes in translation control. Mol Cell Biol 15:2719–27.

68. White J, Li Z, Sardana R, Bujnicki JM, Marcotte EM, Johnson AW. 2008. Bud23 methylates G1575 of 18S rRNA and is required for efficient nuclear export of pre-40S subunits. Mol Cell Biol 28:3151–61.

69. Sikorski RS, Hieter P. 1989. A system of shuttle vectors and yeast host strains designed for efficient manipulation of DNA in Saccharomyces cerevisiae. Genetics 122:19–27.

